# Intrarenal B cells integrate *in situ* innate and adaptive immunity in human renal allograft rejection

**DOI:** 10.1101/2020.09.26.314377

**Authors:** Yuta Asano, Joe Daccache, Dharmendra Jain, Kichul Ko, Andrew Kinloch, Margaret Veselits, Donald Wolfgeher, Anthony Chang, Michelle Josephson, Patrick Cunningham, Anat Tambur, Aly Khan, Shiv Pillai, Anita S. Chong, Marcus R. Clark

## Abstract

In human allograft rejection, intrarenal B cell infiltrates identify those with a poor prognosis. However, how intrarenal B cells contribute to rejection is not known. Single cell RNA-sequencing of intrarenal class-switched B cells revealed a unique innate cell transcriptional state resembling murine peritoneal B1 cells (Bin cells). Comparison to the transcriptome of whole renal allograft rejecting tissue revealed that Bin cells existed within a complex autocrine and paracrine network of signaling axes. The immunoglobulins expressed by Bin cells did not bind donor specific antigens nor were they enriched for reactivity to ubiquitously expressed self-antigens. Rather, Bin cells frequently expressed antibodies reactive with renal expressed antigens. Furthermore, local antigens could drive Bin cell proliferation and differentiation into plasma cells expressing self-reactive antibodies. By contributing to local innate immune networks, and expressing antibodies reactive with renal expressed antigens, Bin cells are predicted to amplify local inflammation and tissue destruction.

## Introduction

In germinal center (GCs), spatial and molecular orchestration of clonal expansion, somatic hypermutation (SHM) and selection drive production of high affinity antibodies and immunological memory *(Kennedy et al., 2020; Victora and Nussenzweig, 2012)*. Remarkably, in many inflammatory and autoimmune diseases, GC-like structures form in afflicted organs (tertiary lymphoid structures, TLS) *(Leslie, 2016)*. Furthermore, TLS are often associated with hallmarks of B cell selection including local clonal expansion and SHM. However, in most human diseases, the antigens driving *in situ* selection are not known. We do not know if TLS-mediated selection usually results in high-affinity and specific antibodies. Furthermore, it is not clear if the GC B cell molecular programs are recapitulated in TLS *(Kennedy et al., 2020)*. Finally, we do not know if B cells can be selected in inflamed tissue in the absence of histologically obvious TLS. Therefore, our fundamental understanding of local *in situ* adaptive immunity in human disease is incomplete.

Acute renal allograft rejection is associated with B cell infiltrates *(Alsughayyir et al., 2017)* which, in most studies, predict poor graft survival *(Hippen et al., 2005; Sarwal et al., 2003; Tsai et al., 2006)*. B cells are often organized into TLS and, in mice *(Tse et al., 2015)* and humans *(Steinmetz et al., 2007)*, B cell depletion appears to mitigate rejection. There are myriad mechanisms by which infiltrating B cells could be pathogenic including secreting antibodies, inflammatory cytokines and antigen presentation. Alternatively, it is possible that the majority of infiltrating lymphocytes are non-specifically trapped in inflamed renal tissue *(Walch et al., 2013)*. In this scenario, they would merely be a bystander of other, central pathogenic processes. Discriminating between these possibilities requires understanding the immunoglobulin repertoire and phenotype of infiltrating B cells in acute allograft rejection.

One clear pathogenic function of B cells in acute allograft rejection is the secretion of donor human leukocyte antigen (HLA)-specific antibodies (DSA). Serum DSA strongly predict early onset of allograft rejection *(Djamali et al., 2014; Lawrence et al., 2013; Zhang, 2018)*. The source of these DSA is not known. A study of a single infected end-stage kidney explant suggested infiltrating B cell infrequently expressed DSAs *(Porcheray et al., 2012)*. In another analysis, two antibodies, whose heavy and light immunoglobulin chains were paired based on bulk renal allograft RNA-seq, suggested *in situ* selection for lipopolysaccharide (LPS) reactivity *(Grover et al., 2012)*. Therefore, from these limited studies, it is unclear if infiltrating B cells express DSAs during ongoing rejection.

Analysis of renal bulk RNA has suggested clonal expansion of B cells expressing somatically hypermutated antibodies *(Cheng et al., 2011; Ferdman et al., 2014)*. However, it is unclear if the observed clonal expansion was a general feature of infiltrating B cells or if clonality arose from infrequent plasma cells. Indeed, the latter possibility is consistent with the association of infiltrating plasma cells, and not CD20^+^ B cells, with serum DSAs. In both mice and humans, renal transplant rejection can be associated with loss of tolerance and serum antibodies to self-antigens *(Cardinal et al., 2017)*. However, the source of these autoantibodies is also not known. Therefore, in ongoing renal allograft rejection, the antigens driving *in situ* B cell selection, and even the magnitude of that selection, remain unknown.

In addition to antibody production, B cells modulate inflammation by presenting antigens to T cells *(Liarski et al., 2014)* and secreting cytokines. Of potential relevance to renal allograft tolerance are regulatory B (Breg) cells secreting IL-10 *(Laguna-Goya et al., 2020; Peng et al., 2018)*. Tolerant renal allograft recipients have increased circulating B cells *(Newell et al., 2010; Pallier et al., 2010)*. Moreover, in one small study pan-B cell depletion by rituximab paradoxically induced acute allograft rejection *(Clatworthy et al., 2009)*. Therefore, different B cell populations, in different contexts, might play opposing roles in allograft rejection. We do not know the phenotype and heterogeneity of intrarenal B cell populations and therefore cannot predict whether they mediate or repress ongoing rejection. There are large and critical gaps in our knowledge of the phenotype and function of intrarenal B cells in renal allograft rejection.

Herein, using single cell RNA sequencing (scRNA-seq) we report that in allograft rejection, intrarenal B cells have a unique transcriptional state that resembles murine B1 innate-like B cells. Such cells have not been previously identified in humans. Bin cells are not a source of DSAs. Rather, they can express renal-reactive antibodies and give rise to proliferating plasma cells selected by local antigens. These results demonstrate how intrarenal B cells drive local inflammation and contribute to allograft rejection.

## Results

### Distinct transcriptional states in activated intrarenal and tonsil B cells

To better understand *in situ* B cell responses in renal allograft rejection, we sorted CD45^+^ DAPI^-^ Calcein^+^ CD19^+^ CD38^+^ activated B cells from five renal allograft patient biopsies and four tonsillectomy patient samples (Figure 1A). A paired biopsy from each renal allograft patient was reviewed by a blinded renal pathologist. Presence of B cells and C4d deposition was examined by immunohistochemistry and serum assayed for DSA. Results and clinical characteristics for each patient are provided in Supplementary Table 1.

**Figure 1.**
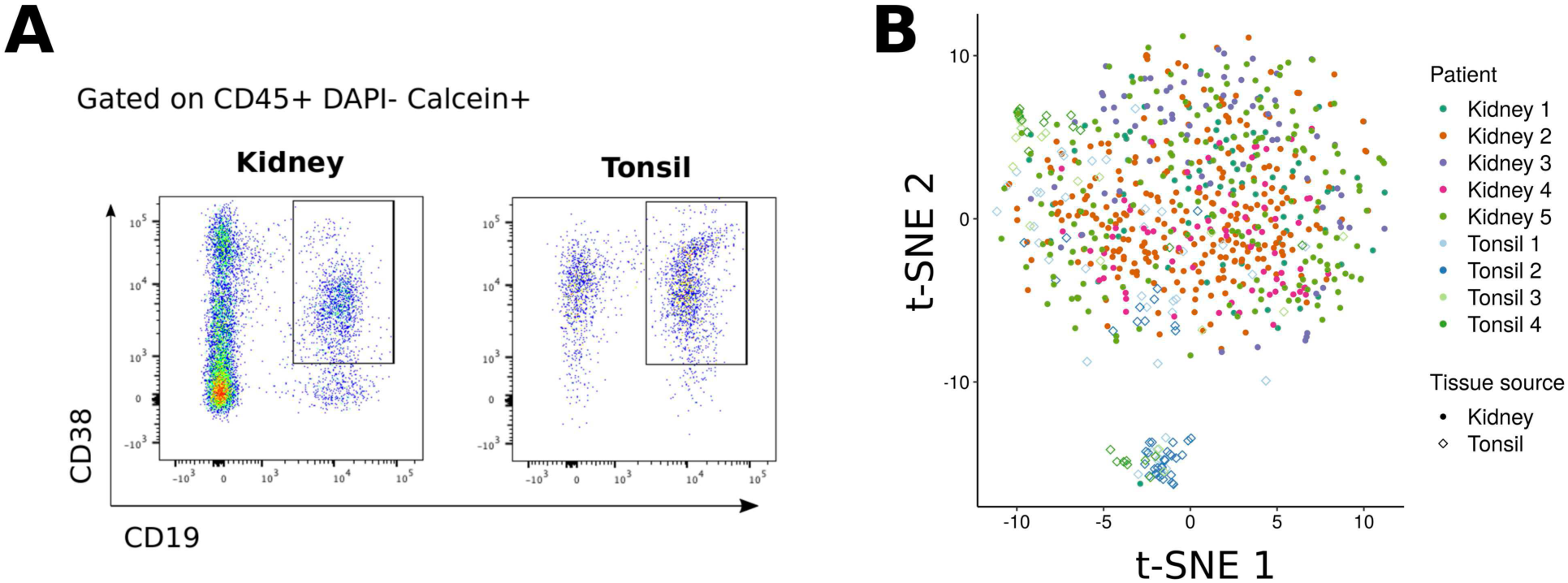
Sorting and scRNA-seq of activated B cells in rejecting renal allograft and tonsil. (A) Gating scheme for single-cell sorting of CD19^+^ CD38^+^ activated B cells in renal allograft and tonsil samples. (B) A t-SNE plot of scRNA-seq. Color and shape respectively indicate patients and tissue sources from which cells were derived.

Sorted B cells were then subjected to scRNA-seq following the Smart-Seq2 protocol *(Picelli et al., 2014)*. To ensure a high-quality dataset, we excluded cells which had less than 3,000 or more than 15,000 expressed genes (Supplementary Figure 1A). Additionally, to exclude non-B cells, we removed cells which had low expression of immunoglobulin (Ig) constant region genes. After the quality control (QC), 655 renal and 129 tonsil B cells were used for subsequent analyses. Batch effects from separate sequencing runs were normalized using External RNA Control Consortium (ERCC) spike-in control and RUVSeq R package *(Risso et al., 2014)* (Supplementary Figure 1B and 1C).

To assess cell population heterogeneity, we mapped sequenced cells onto a t-distributed stochastic neighbor embedding (t-SNE) space. As demonstrated in Figure 1B, renal B cells formed one diffuse cluster while tonsil B cells formed two distinct clusters, one of which overlapped with the kidney cluster and the other that was distinct (Figure 1B). This clustering was not due to batch-associated differences, suggesting that B cells in renal allograft and tonsil had distinct transcriptional profiles. Moreover, B cells from all five renal biopsies were similarly distributed in the t-SNE space suggesting that renal allograft-infiltrating B cells had a similar transcriptional profile across patients regardless of their histological or clinical features.

Although intrarenal B cells formed a single diffuse cluster on t-SNE, this population could be divided based on Ig class switching (Figure 2A). Here, B cells expressing IgM or IgD as the most highly expressed Ig isotype were categorized as “unswitched”, and those expressing either IgG, IgA or IgE as “switched.” Unswitched cells composed about 70 % of both renal and tonsil B cells, and the remaining class-switched cells mostly expressed IgG or IgA (Supplementary Table 2). Regardless of Ig class, switched cells were similarly distributed in the t-SNE space (Figure 2B and 2C). The two tonsil B cell clusters were also distinguished by their Ig class. Unswitched tonsil B cells largely overlapped with unswitched renal B cells whereas switched tonsil cells formed a distinct cluster. This clustering differences persisted when Ig constant region genes were removed (Supplementary Figure 2A). These data suggest that B cells in rejected renal allograft and tonsil tissue are similar prior to class-switch recombination (CSR) but diverge thereafter.

**Figure 2.**
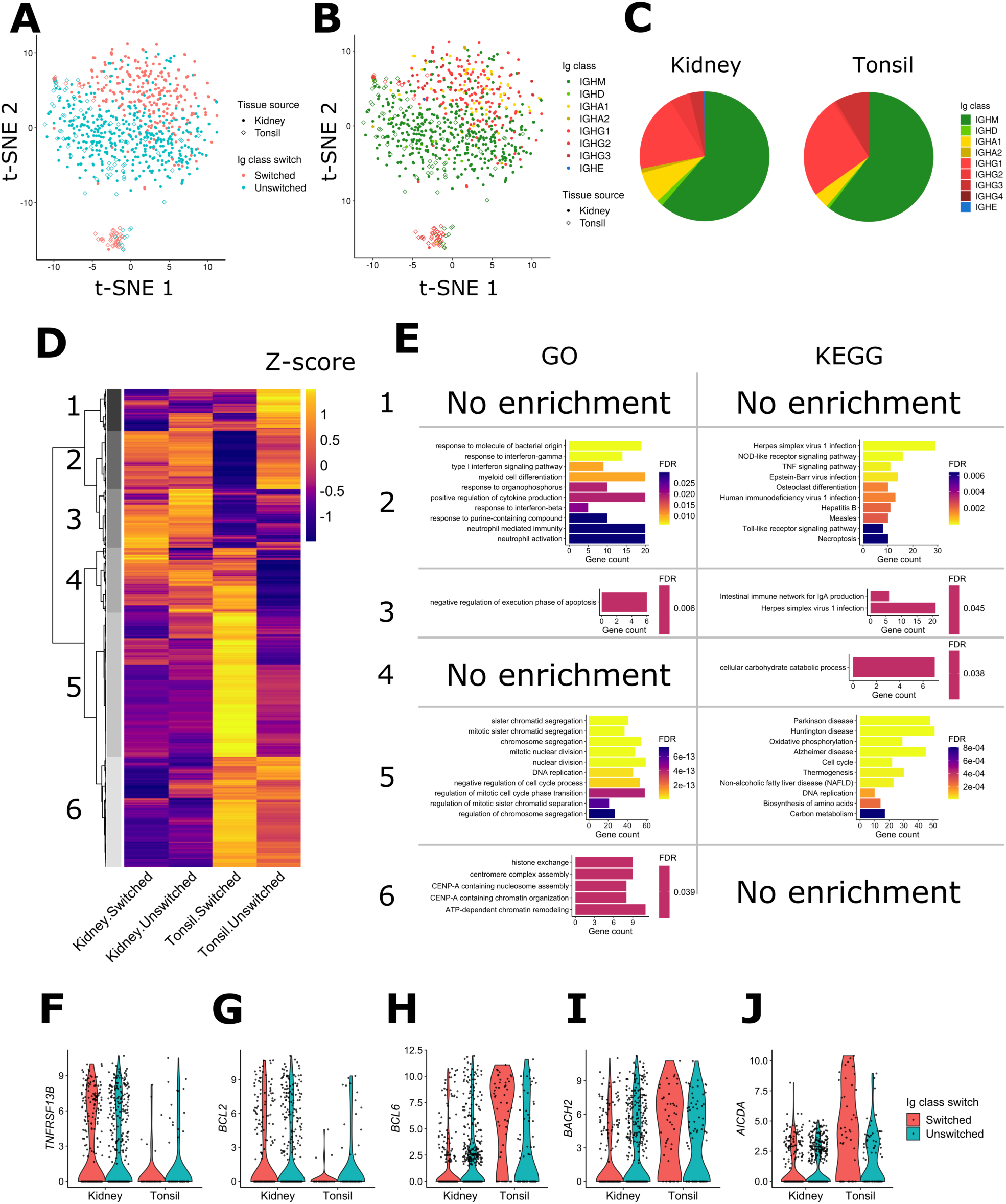
Unique transcriptional state of class-switched intrarenal B cells. (A and B) t-SNE plots as in Figure 1B. Color indicates Ig class-switch state (A) or expressed Ig classes (B). The cells were categorized as “switched” if their most highly expressed Ig heavy chain genes were either IgG, A or E, and categorized as “unswitched” otherwise. (C) Pie charts showing the distribution of Ig class in intrarenal and tonsil B cells. (D) A heatmap showing hierarchical clustering of the 2855 DEGs. Mean expression values were calculated for each four cell populations based on their tissue source and Ig class-switch state and then converted to Z-scores. (E) Enrichment of GO terms and KEGG pathways in the five gene clusters. At most 10 most significantly enriched pathways were shown per cluster. (F-J) Violin plots showing RNA expression of *TNFRSF13B* (F), *BCL2* (G), *BCL6* (H), *BACH2* (I), and *AICDA* (J).

We next compared differential B cell gene expression across tissue sources and Ig class-switch states. This comparison identified 2855 differentially expressed genes which could be divided into six hierarchical clusters (Figure 2D and Supplementary Table 3). Cluster 1 included genes enriched in unswitched tonsil B cells, clusters 2 and 3 genes enriched in intrarenal cells, cluster 4 genes enriched in intrarenal and tonsil switched cells, cluster 5 genes enriched in tonsil switched cells and cluster 6 genes enriched in tonsil B cells. A pathway enrichment analysis based on Gene Ontology (GO) and Kyoto Encyclopedia of Genes and Genomes (KEGG) databases revealed biological pathways enriched in most clusters (Figure 2E).

Many of the GO and KEGG pathways enriched in cluster 2 were related to innate receptors and signaling pathways. Indeed, intrarenal B cells highly expressed specific pattern recognition receptors (PRRs) in clusters 2 and 3 including *NLRP1*, *NOD1*, *TLR2 and TLR7* (Supplementary Table 3). Therefore, we next examined if, globally, clusters 2 and 3 were enriched for innate immune genes. We identified genes tagged to a GO term “innate immune response”. When we calculated a sum of scaled expression values of such genes in cluster 2 and 3, intrarenal cells had higher values than tonsil, especially class-switched, B cells (Supplementary Figure 2B). This enrichment of innate immune response genes was consistent across all patients (Supplementary Figure 2C). Therefore, there is an enrichment for innate immune response genes in intrarenal B cells.

Cluster 2 was also enriched in interferon (IFN)-related pathways. IFN-γ signaling induces NLRC5, an important MHC class-I activator *(Kobayashi and van den Elsen, 2012)* and one of the upregulated innate receptors in cluster 3. In addition, these clusters contained genes encoding a type-1 IFN receptor IFNAR2, along with signaling molecules shared between type-1 and type-2 IFN pathways such as IRF1 and STAT1. These results suggest an overall activation of IFN pathways in intrarenal B cells.

Clusters 2 and 3 also contained several cytokine ligands and receptors: *IFNAR2, IL15*, *TNFRSF1B*, and *TNFRSF13B* (Figure 2F). *TNFR13B* encodes TACI, a receptor for BAFF. BAFF mediates B cell survival and is important in some B-cell-mediated inflammatory diseases *(Davidson, 2010)*. Furthermore, BAFF over-expression is associated with renal allograft rejection *(Pongpirul et al., 2018; Wang et al., 2019)*. Therefore, the high expression of TACI in intrarenal B cells suggests a role in *in situ* activation and survival.

The anti-apoptotic factor *BCL2* was also found in cluster 2 (Figure 2G) which is consistent with our previous report of high *BCL2* expression in intrarenal B cells in human renal allograft rejection and lupus nephritis *(Ko et al., 2016)*. Along with *BCL2*, many of the pathways enriched in cluster 2, such as interferon, Toll-like receptor (TLR), or cytokine-production-related pathways are direct targets of BCL6, a transcriptional repressor in GC responses *(Basso et al., 2010)*. Indeed, expression of *BCL6* was lower in renal B cells (Figure 2H), as well as another transcriptional repressor *BACH2*, which shares targets with BCL6 *(Huang et al., 2014)* (Figure 2I). These results suggest that some of the intrarenal B cell phenotype might be due to reduced *BCL6* and *BACH2* expression.

*BCL6* and *BACH2* were found in cluster 5 and 6 respectively, which contained genes upregulated only in class-switched tonsil B cells. These cells were enriched in several pathways that have previously been ascribed to GC B cells including proliferation and somatic hypermutation. Notably, *AICDA* was expressed in class-switched tonsil B cells but not significantly in other B cell populations (Figure 2J). These results indicate that intrarenal class-switched B cells lack the essential transcriptional features of GC B cells.

Cluster 3 captured genes specifically upregulated in intrarenal B cell populations while cluster 4 genes were preferentially expressed both in intrarenal and class-switched tonsil B cells. Neither gene cluster demonstrated upregulation of specific GO pathways. However, examination of individual differentially expressed genes reveal potentially important differences. Most notable was *AHNAK* (Figure 3A). *AHNAK* mRNA levels were far higher in intrarenal B cells compared to tonsil regardless of Ig class switch (Figure 3B). Expression of the AHNAK protein in intrarenal B cells was confirmed by confocal microscopy of renal allograft rejection and tonsil tissue (Figure 3C). Although AHNAK has been reported to be important for regulation of calcium signaling in activated T cells *(Matza et al., 2008)*, its role in B cell biology is unknown. Interestingly, within mouse B cell subsets, *Ahnak* is preferentially expressed in peritoneal cavity B1a and B1b cells (Immgen, Figure 3D) *(Heng and Painter, 2008)*. Furthermore, this expression pattern was shared with murine homologues of several other cluster 3 genes, such as *ITGAM* and *VIM* (Supplementary Figure 2D and 2E). Therefore, we examined whether cluster 3 was enriched for genes having an *AHNAK* covariant expression pattern.

**Figure 3.**
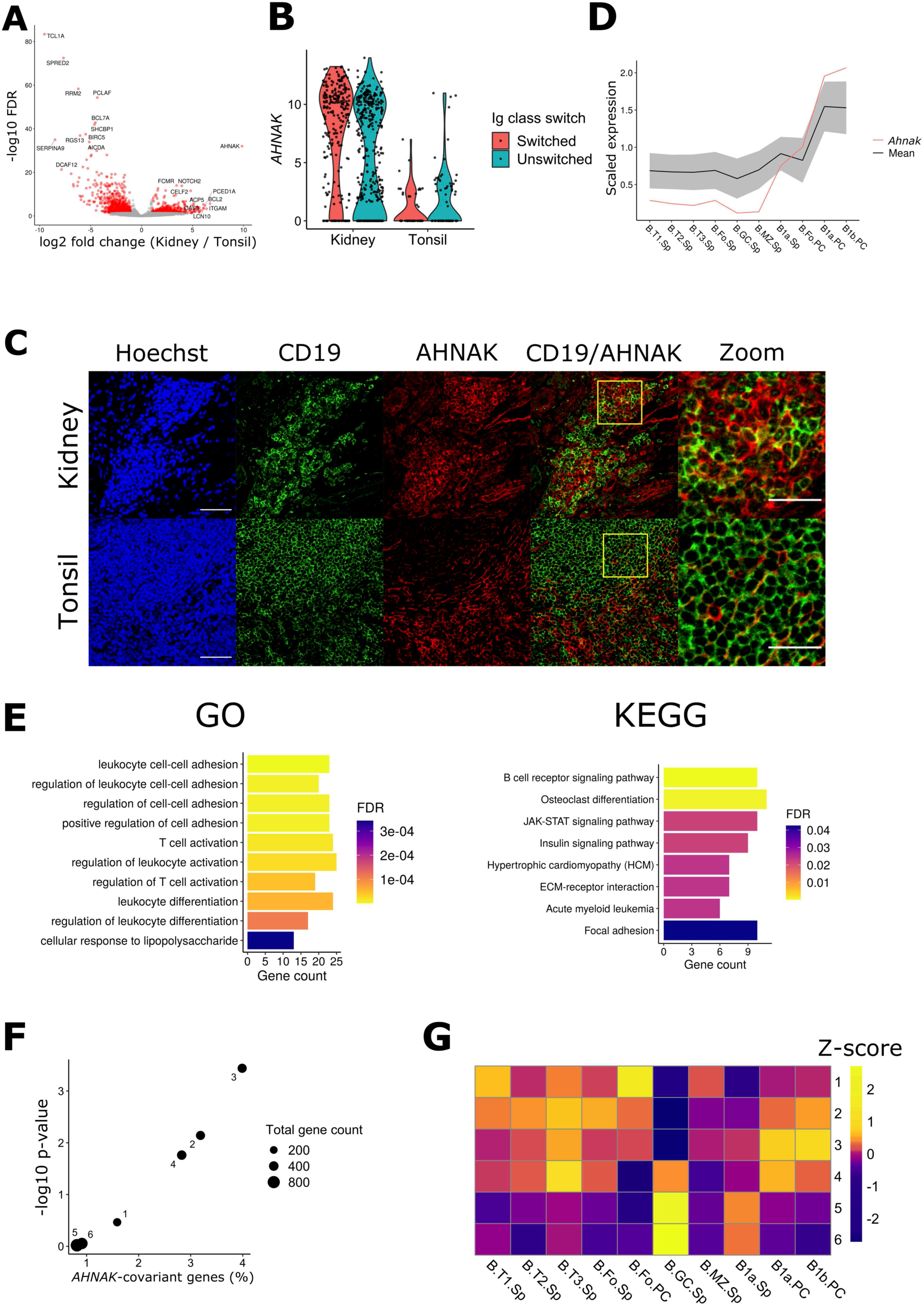
Intrarenal B cells have an innate-like gene signature. (A) A volcano plot showing DEG between Ig class-switched intrarenal and tonsil B cells. Genes expressed higher in intrarenal B cells are shown on the right side of the plot. (B) A violin plot demonstrating RNA expression of *AHNAK*. (C) Staining images of AHNAK with nuclei (Hoechst) and a B-cell marker CD19 in rejected renal allograft and tonsil. The high-magnification panel corresponds to the yellow square on the merged panel. Scale bars indicate 50 μm or 25 μm (high-magnification panel). (D) Expression of 333 murine genes that correlate with *Ahnak* in Immgen. The mean value of the 333 *Ahnak*-covariant genes (including *Ahnak* itself) is shown as the black line with the grey shade indicating standard deviation. Expression of *Ahnak* is the red line. T: transitional, Fo: follicular, GC: germinal center, MZ: marginal zone, Sp: spleen, and PC: peritoneal cavity. (E) Enrichment of GO terms and KEGG pathways in the 293 AHNAK-covariant genes. At most 10 most significantly enriched pathways are shown. (F) Enrichment of the AHNAK-covariant genes in each gene cluster from Figure 2D. (G) A heatmap showing DGE scores, a sum of scaled expression levels of each gene cluster within each murine B cell subset in Immgen data. Each row and column represents the gene clusters found in Figure 2D and the murine B cell subpopulations. DEG scores were scaled by row to obtain Z-scores.

We identified 333 mouse genes whose expression pattern in peripheral B cell populations was similar to *Ahnak* (correlation coefficient ≥ 0.8) (Figure 3D and Supplementary Table 4). The *Ahnak-*covariant murine genes corresponded to 293 human homologues. These human genes were enriched for cell adhesion and lymphocyte activation pathways, as well as an innate-immune pathway related to lipopolysaccharide responses (Figure 3E). The *AHNAK-*covariant genes were highly enriched in cluster 3 and to a lesser degree in clusters 2 and 4 (Figure 3F). These results suggest that *AHNAK*-covariant genes are a signature of intra-graft B-cell activation.

Although the *AHNAK*-covariant genes were enriched in cluster 3, they represented only 5% of the cluster. Therefore, we next tested if cluster 3 genes were generally enriched in peritoneal cavity B1 cells. First, we converted the 2855 differentially expressed genes into murine orthologs. Then, in the Immgen data, we compared murine B cell subsets for expression of genes in each human gene cluster. For each gene cluster, we calculated a sum of gene expression values scaled across the B cell subsets to evaluate overall gene expression. This analysis demonstrated that genes in cluster 3 were most highly expressed by peritoneal cavity B1 cells (Figure 3G). This pattern in Z-scores was weaker, but still present, when scores were calculated without the *Ahnak*-covariant genes (Supplementary Figure 2F). This result suggests that cluster 3, containing genes highly expressed in the graft-infiltrating B cells, have a gene signature of peritoneal B1 cells.

We also investigated if intra-graft B cells expressed gene programs previously ascribed to infiltrating B cells associated with other inflammatory renal diseases. One such pathway is the age-associated B cell (ABC) or double negative (CD27^-^IgD^-^, DN) pathway. DN-like cells have been identified in tissue inflammation including lupus nephritis *(Arazi et al., 2019)*, and are characterized by increased expression of *TBX21* (T-bet), *ITGAX* (CD11c), *TLR2* and *TLR7*. Although both *TLR2* and *TLR7* are preferentially expressed by intra-graft B cells, we found no difference between intrarenal and tonsil B cells in the overall expression of 26 genes associated with DN cells *(Arazi et al., 2019; Jenks et al., 2018)* (Supplementary Figure 3A). Therefore, although DN cells and our intrarenal B cells share high expression of innate immune genes, otherwise they have distinct phenotypes.

We also examined the HIF-1 pathway, which is upregulated *in situ* in both human and murine lupus nephritis *(Chen et al., 2020; Yang and Liu, 2016)*. However, HIF-family genes were not upregulated in intrarenal B cells compared with tonsil B cells (Supplementary Figure 3B-3F). These data indicate that the hypoxia pathway is not characteristic for intrarenal B cells in acute allograft rejection. Overall, these data demonstrate that class-switched intrarenal B cells in allograft rejection have a unique transcriptional profile reminiscent of innate B1 cells, which is distinct from intrarenal B cells in lupus nephritis.

### Intrarenal signaling networks

We next examined the x other cell populations in rejecting renal allografts. For this purpose, we first identified genes known or postulated to encode a ligand or receptor *(Ramilowski et al., 2015)* that were preferentially expressed in Ig class-switched renal versus tonsil B cells. We then queried whole tissue microarray data from patients with renal allograft rejection for preferentially expressed ligands and receptors *(Sarwal et al., 2003)*. With these gene sets, we tested if there were ligand-receptor pairs co-upregulated between the intrarenal B cells and rejected tissues.

We identified 10 ligand-receptor pairs coordinately upregulated in B cells and renal tissue (Figure 4A). One of them was a pair of genes encoding the pro-inflammatory cytokine IL-15 (Figure 4B) and a component of its receptor complex, IL-15RA. Tissue staining showed no detectable IL-15 expression in tonsil germinal centers whereas expression was readily detected in infiltrating B and other immune cells in rejected renal allografts (Figure 4C). IL-15RA was moderately expressed in both tonsil and renal graft tissue. However, it was more abundant in renal tubular cells than infiltrating immune cells (Figure 4C), suggesting that IL-15 secreted by B cells might be captured by the tubular cells for presentation to immune cells *(Abadie and Jabri, 2014)*. These data suggest an interplay between intrarenal B cells, tubular cells and other cellular components in allograft rejection.

**Figure 4.**
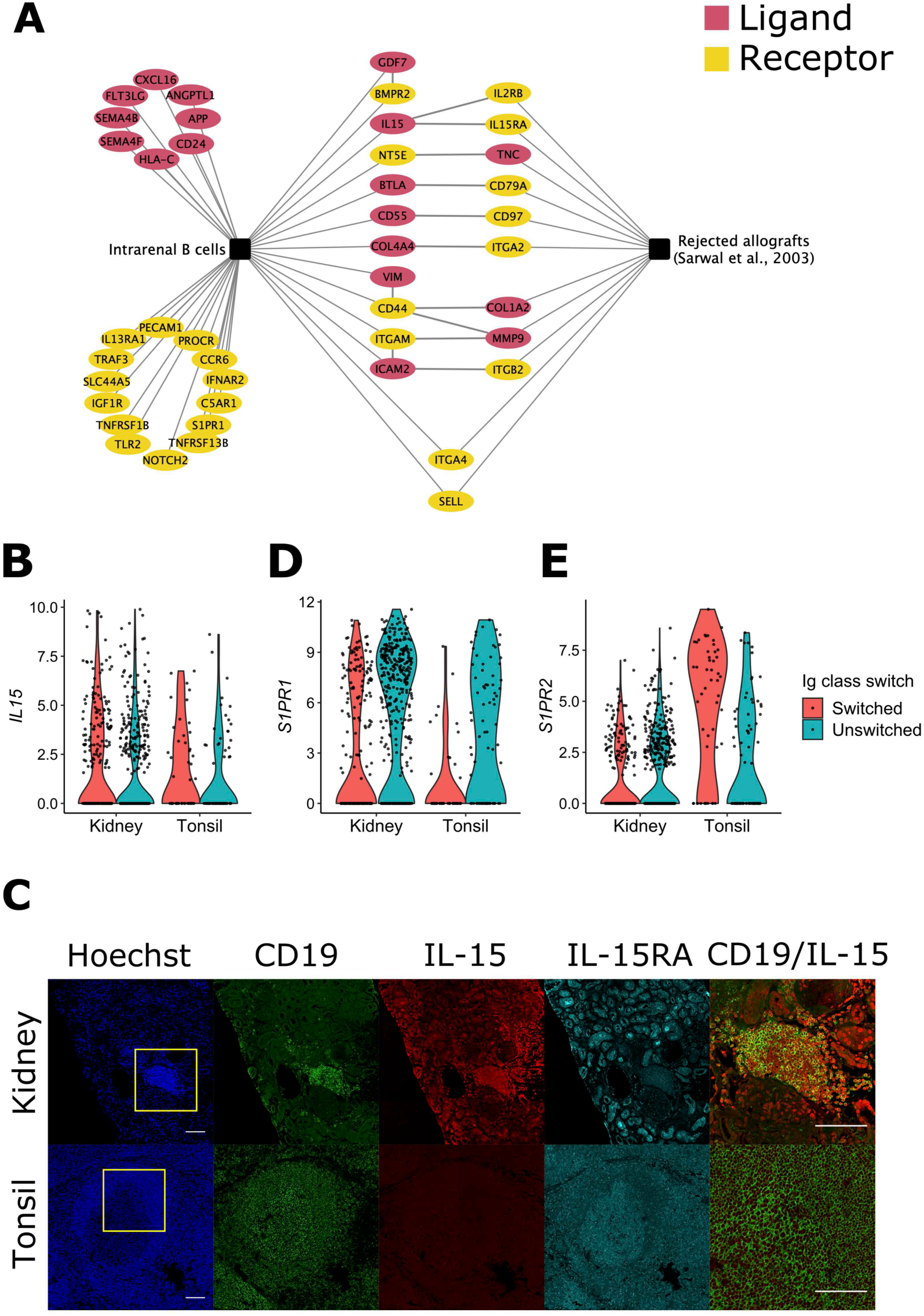
Co-regulated ligand-receptor pairs in intrarenal B cells and renal allograft tissue. (A) Ligand and receptor genes upregulated in Ig-switched renal B cells (compared with Ig-switched tonsil B cells) or genes upregulated in rejected renal allograft tissues (compared with normal allografts). Gene names were connected to each other when both their ligand and receptor were present. Color (red and blue) indicates ligands and receptors respectively. (B) A violin plot showing the expression level of *IL15*. Cells were grouped by their tissue source and color indicates Ig class-switch state. (C) Staining images of nuclei (Hoechst), CD19, IL-15, and IL-15RA in rejected renal allograft and tonsil tissues. The merged CD19/IL-15 panels were a magnification of the yellow box in merge panels. Scale bars indicate 100 μm. (D-E) Violin plots as in (B), showing the expression level of *S1PR1* (D) and *S1PR2* (E).

The other co-expressed pairs contained several cell migration-related axes, such as *CD55*-*CD97*, *COL4A4*-*ITGA2*, *COL1A2*-*CD44*, *ICAM2*-*ITGB2* (Figure 4A). Although not paired with their ligands, both B cells and renal tissue also expressed *ITGA4* and *SELL*. These receptor-ligand pairs provide a potential mechanism for the migration and/or retention of B cells within rejecting renal allografts.

Another co-upregulated pair was *BTLA* and *CD79A*. *CD79A* encodes Igα which is a signaling chain of BCR. Therefore, it is expressed ubiquitously and specifically in B cells. Its ligand BTLA is reported to attenuate BCR signaling through Igα binding on the same cell surface *(Vendel et al., 2009)*. Therefore, the *BTLA-CD79A* axis could modulate *in situ* B cell responses to antigen within rejecting allografts.

Several receptor and ligand pairs were expressed in B cells including, *GDF7*-*BMPR2*, *VIM*-*CD44* and *ITGAM*-*ICAM2*. Co-expression in B cells provide for several potential autocrine signaling axes for homotypic adhesion and activation (Figure 4A). Although not paired with tissue-upregulated genes, intrarenal B cells expressed other migration-related genes: *APP, CD24, CXCL16, CCR6, PECAM1* and *S1PR1* (Figure 4A). S1PR1 is critical for B cell egress from lymphoid organs *(Cyster and Schwab, 2012)*, and is downregulated in GC B cells. Instead, S1PR2 is upregulated in GC B cells and coordinates their migration *(Green and Cyster, 2012)*. Relatively high expression of *S1PR1*, and low expression of *S1PR2* (Figure 4D and 4E), suggest that renal B cells respond to different localization signals than GC B cells. These data indicate that unique receptor-ligand networks regulate local Bin cell retention and activation.

### Serum DSA is not predictive of intrarenal B cell phenotype

In renal allograft rejection, the presence of serum DSA predict a worse clinical outcome *(Djamali et al., 2014; Lawrence et al., 2013; Zhang, 2018)*. Currently, the paradigm is that DSA are produced in secondary lymphoid organs. However, they could arise from intrarenal B cells. To begin to address this possibility, we sought to determine if the presence of serum DSA was associated with differences in the transcriptional programs of intrarenal B cells. Since our initial cohort had only one DSA-positive patient, we obtained three additional renal biopsies (two from DSA-positive patients and one from a DSA-negative patient) as well as three additional tonsillectomy samples. From these samples, activated B cells were sorted and subjected to scRNA-seq (Supplementary Table 1).

After filtering for both gene coverage and Ig expression, we obtained an additional 513 cells to integrate with those of the first cohort (Supplementary Figure 4A). The second cohort had a lower sequence coverage compared to the first cohort (Supplementary Figure 4B). Since normalizing scRNA-seq data with different sequencing depth by a single-scaling factor (e.g. total mapped reads) could introduce bias *(Bacher et al., 2017; Hafemeister and Satija, 2019)*, we first normalized our data by SCTransform implemented in Seurat *(Hafemeister and Satija, 2019; Stuart et al., 2019)*. The data were then integrated by ComBat *(Leek et al., 2012)* to negate batch effects between cohorts (Supplementary Figure 4C and 4D).

The resulting integrated data had a similar t-SNE projection to that of the first cohort reproducing the distinct renal and tonsil B cell clusters (Figure 5A). As in the previous analysis, class switched B cells were separate whereas unswitched cells clustered together (Figure 5B). In addition, there was a new discrete cluster which highly expressed *PRDM1* indicating they were plasma cells (Figure 5C and 5D). This population was mostly derived from kidney patient 6 and, to a lesser degree, patient 7. A few plasma cells were also detected in biopsies from patients 2, 5 and 8. Notably, plasma cells were observed in biopsies from two DSA-positive and three DSA-negative patients. Therefore, intra-renal plasma cells were not a unique feature of DSA-positive patients.

**Figure 5.**
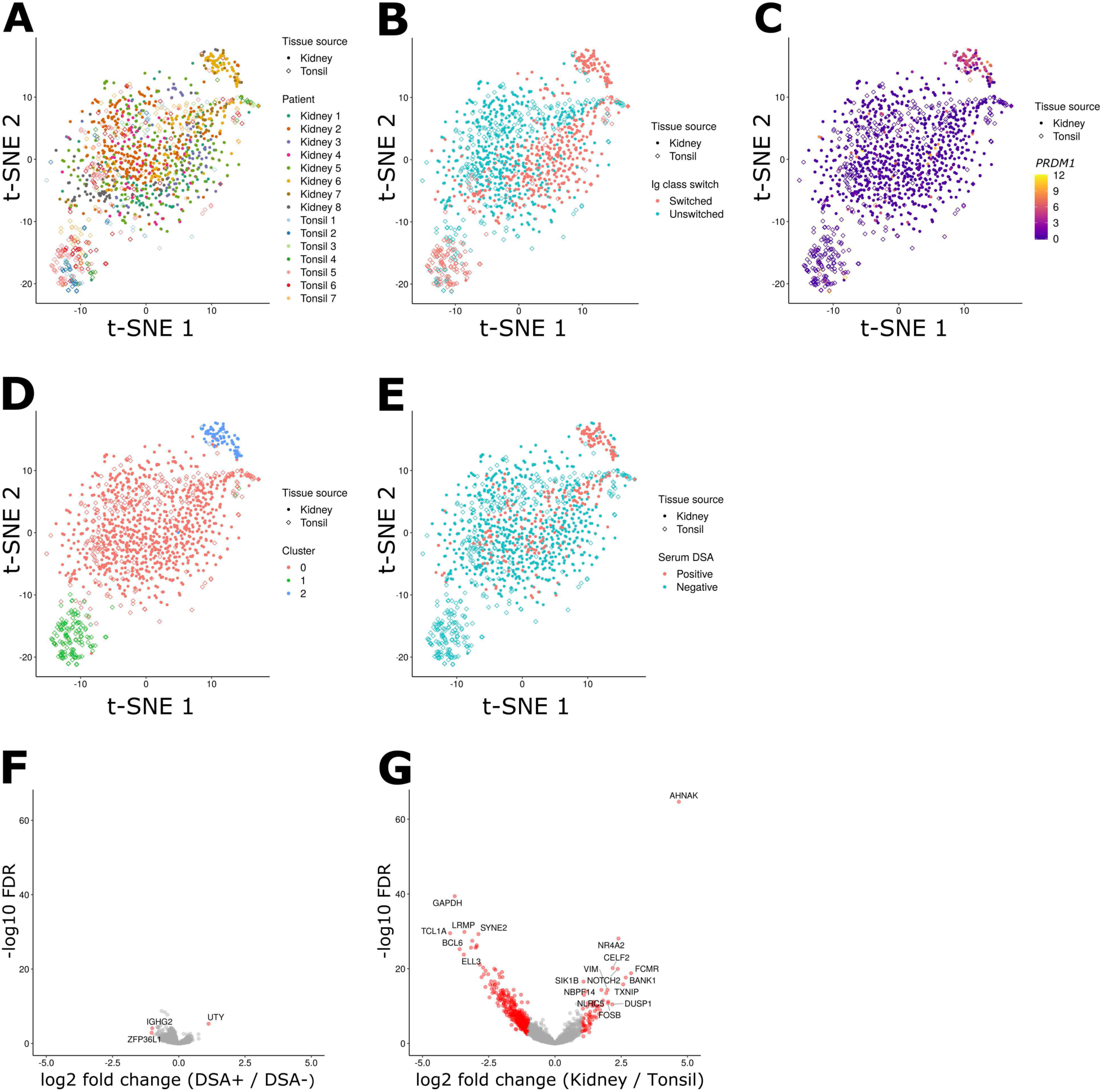
B cells from serum DSA-positive and negative patients are similar. (A-E) t-SNE plots of the integrated data of the two cohorts. Shape indicates tissue source, and color indicates patients (A), Ig class-switch state (B), *PRDM1* expression (C), clusters assigned by Seurat (D), and serum DSA positivity (E). (F and G) Volcano plots showing differential gene expression in non-plasma B cells, comparing DSA-positive and negative patients (E), or Ig class-switched renal and tonsil cells (F). Shown on the right side of the plots are genes upregulated in B cells from DSA-positive patients (F) or intrarenal B cells (G). Genes above the significance threshold were colored in red, and names of top-hit genes were labeled.

The other B cell populations did not cluster depending on patients’ serum DSA positivity (Figure 5E). Comparing gene expression of the non-plasma cells from the DSA-positive and DSA-negative patients, we could identify only 3 genes differentially expressed (Figure 5F). In contrast, we detected nearly 500 differentially expressed genes between class-switched renal and tonsil non-plasma cells in this combined dataset (Figure 5G). These results suggest that there are no substantial transcriptional differences in graft-infiltrating B cells from DSA-positive and DSA-negative patients.

### In situ immunoglobulin repertoire in renal allograft rejection

A central question is whether intra-graft B cells are selected for allo reactivity. A previous study of one renal explant suggested that graft-infiltrating B cells infrequently expressed HLA-reactive antibodies and none of these were donor specific*(Porcheray et al., 2012)*. However, this study did not address whether B cells are selected *in situ* for HLA reactivity in ongoing renal transplant rejection.

Therefore, we first used nested polymerase chain reaction (PCR) to amplify immunoglobulin gene variable regions from the cDNA of single B cells isolated from seven kidney biopsies and two tonsil samples as previously described *(Smith et al., 2009)*. Obtained sequences were aligned to IMGT reference using IMGT/HighV-QUEST *(Alamyar et al.; Brochet et al., 2008)* and analyzed for somatic mutations. In total, we identified full-length Ig heavy chain variable regions from 457 B cells in seven rejection patients and 77 in two tonsillectomy patients (Figure 6A). Overall, immunoglobulin mutation burden was similar between rejection and tonsillectomy samples (Figure 6B). This was true for both B cells expressing unswitched and switched immunoglobulin genes. In general, unswitched B cells had less of a mutation burden than switched cells. However, in intrarenal B cells there were a small fraction of both unswitched and switched cells that had very high (>60) frequencies of mutations.

**Figure 6.**
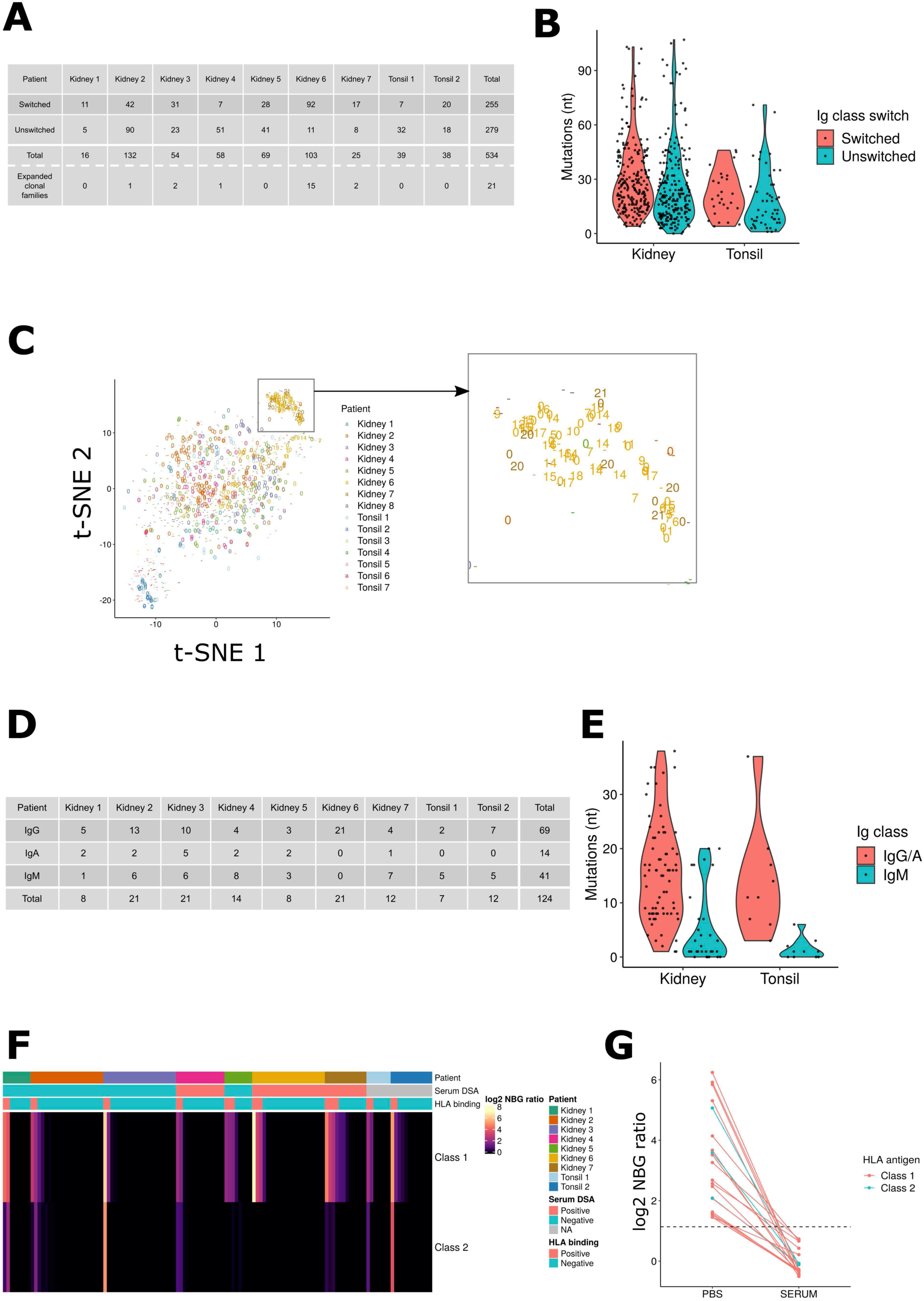
Antibodies expressed by intrarenal B cells were not selected for allo-HLA reactivity. (A) A table showing distribution of number and Ig class of the sequenced antibody heavy chains across the patients. Number of expanded clonal families per patient is also shown. (B) A violin plot showing distribution of mutations in the variable region of immunoglobulin heavy chains, grouped by tissue source and the Ig class-switch state. (C) t-SNE plots as in Figure 6, with data points labeled with their clonal family. "-" means antibody genes were not sequenced; "0" means the cells did not share their clonotype with others; and 1-21 indicate that the cells shared a clonotype with other B cells with the same number. (D) A table showing distribution of number and Ig class of the recombinantly produced antibodies across the patients. (E) A violin plot showing distribution of mutations in recombinantly expressed antibodies, grouped by tissue source and Ig class-switch state. (F) A heatmap showing the HLA-binding assay result. Each column represents each antibody, and maximum NBG ratio within class-1 or class-2 beads are shown. The header label indicates patients from whom the antibodies were derived, serum DSA positivity, and whether the observed reactivity was above the positivity threshold. (G) A plot showing the change in HLA binding when the antibodies were tested in negative control human serum. All the positive intrarenal antibodies in Figure 6F were tested, and data points of the same antibodies were connected by lines.

Next, we investigated clonal relationships among the sequenced antibodies. We tested whether the B cells were clonally related, defined as sharing the same variable (V), diversity (D), joining (J) segments and complementarity-determining region (CDR) 3 length. Despite high mutation rates, we found only a limited number of shared clonal families in most patients (Figure 6C). In contrast, many of the plasma cells were clonally related, comprising 15 clonal families in patient 6, and 2 clonal families in patient 7. This suggests that plasma cell populations were clonally expanded, and raised the possibility that they were selected *in situ* for specific antigens. In contrast, the lack of clonality of intrarenal B cells suggest that they may have been recruited as highly mutated B cells from peripheral blood.

Many of the clones identified in plasma cells were also present in the corresponding B cell populations. This was true for eight clones in patient 6 and one clone in patient 7. These data indicate that *in situ* B cells can be selected by antigen to locally differentiate and then clonally expand.

In order to characterize their immuno-reactivity, we next expressed the cloned BCR genes as recombinant antibodies. Cloned paired heavy and light Ig variable regions were cloned into human IgG expression vectors with the C-terminal of the heavy-chain construct was modified to include a FLAG tag *(Kinloch et al., 2020)*. Antibodies were expressed in Hek293A cells and purified with protein A beads. In total, we expressed 105 antibodies (74 IgG/A and 31 IgM) from intrarenal B cells isolated from 7 patients (Figure 6D, Supplementary Table 5). This included 11 out of the 21 expanded BCR clones from plasma cells expressing IgG. We also expressed 19 antibodies from the two tonsillectomy patients (9 IgG and 10 IgM). As expected, the mutation rate was higher in IgG/IgA tonsil antibodies than IgM (Figure 6E).

### In situ B cells are not selected for alloreactivity

We first tested reactivity of *in situ* expressed antibodies to HLA antigens using a Luminex-based assay in which beads were coated with a mixture of HLA class-I or class-II antigens *(Picascia et al., 2012)*. Antibody binding was evaluated by the normalized background (NBG) ratio, which is fold-increase binding over negative control serum. Binding was positive if it was equal to or higher than 2.2. When diluted in phosphate-buffered saline (PBS), 17% (18/105) bound to screening HLA beads (Figure 6F). In serum DSA-positive patients, 9 of 47 (19%) expressed monoclonal antibodies (mAbs) were HLA-binding, while 16% (9/58) of mAbs from DSA-negative patients bound HLA (p = 0.82, chi-squared test). However, we observed 21% (4/19) positivity in tonsil antibodies. Collectively, these data suggested that HLA-reactivity was not a specific feature of *in situ* expressed antibodies from DSA-positive patients.

To further delineate the specificity of HLA binding, mAbs that bound to the screening HLA beads were tested on single HLA-antigen beads (SAB). Out of 18 tested antibodies, 17 antibodies showed trimmed mean fluorescent intensity (MFI) higher than 1,000. Unexpectedly, 15 of the 17 antibodies bound to HLA-C, with HLA-Cw*06:02 as the top hit (Supplementary Figure 5A and Supplementary Table 6). Furthermore, 16 of 17 antibodies bound to multiple HLA antigens. These HLA alleles recognized by the mAbs were not donor specific HLAs. Indeed, except for 6-2D3, all bound most strongly to recipient-expressed HLAs (Supplementary Figure 5B). These results suggest that intrarenal B cells often express HLA binding antibodies. However, DSA were rare.

We next examined whether shared eplets (discrete epitopes) *(Duquesnoy, 2006)* between multiple HLA alleles could explain the unexpected broad HLA specificity. However, 6-2D3 mAb had reactivity with several HLA-A and C antigens (Supplementary Figure 5C) that did not share any common eplets. Indeed, majority of mAbs (11 out of 17) that bound multiple HLA Class I alleles did not share an eplet among all their top-10 hits (Supplementary Figure 5D). For the remaining 6 mAbs, the top-10 hits did share an eplet for bound HLA-C antigens (Supplementary Figure 5E and F) *(Tumer et al., 2013; Visentin et al., 2019; Xu et al., 2012)*. Thus, shared eplets could not explain the broad reactivity for the majority of the mAbs.

Another possibility is that the broad reactivity of our mAbs represents low affinity polyreactivity. If so, then adding a non-specific blocking reagent should abrogate binding *(Andrews et al., 2015)*. Therefore, we next retested binding of the above antibodies to the screening HLA beads in presence of negative control human serum. Strikingly, in the presence of serum, all antibodies lost their binding to HLA antigens (Figure 6G). In toto, our results indicate that allo- reactive antibodies are not commonly selected *in situ* during acute renal allograft rejection.

Polyreactivity can be associated with autoreactivity *(Dimitrov et al., 2013)*. Therefore, we assayed binding of the mAbs to human epithelial type-2 (HEp-2) cells by immunofluorescence microscopy. For the mAbs generated from non-plasma Bin cells, the frequency of HEp-2-reactive clones was 18 % (15/84) (Figure 7A), which is similar to that reported for naive or tonsil GC repertoires *(Liu et al., 2018; Wardemann et al., 2003).* These results suggest these antibodies are not selected for autoreactivity.

**Figure 7.**
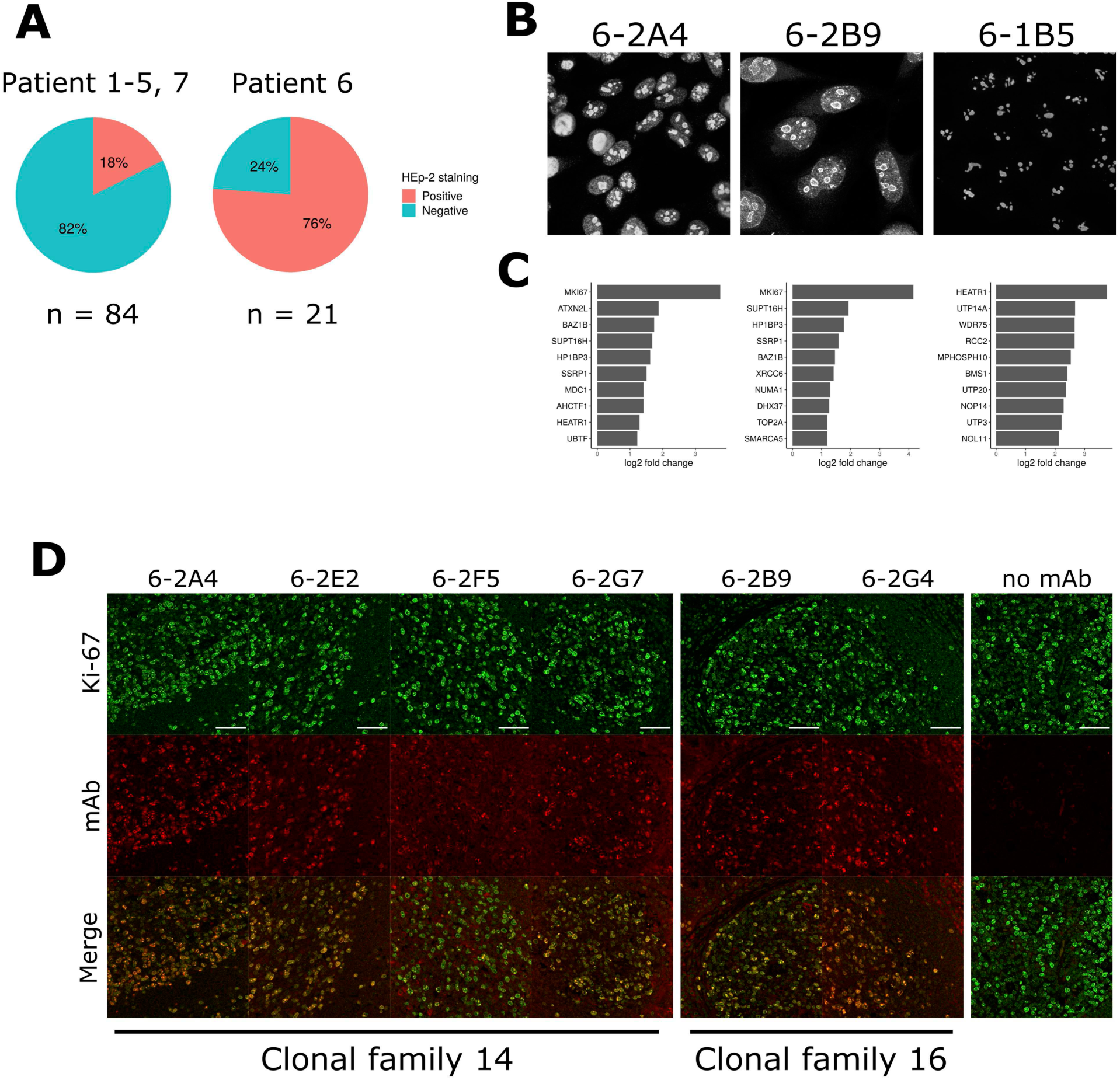
Plasma cells express antibodies specific for nucleolar proteins including Ki-67. (A) Pie charts showing the frequency of HEp-2-reactive antibodies in indicated patients. (B and C) HEp-2 staining images of the three antibodies used for the IP/mass spectrometry (B) and their top 10 preferentially bound antigens (C). Log2 fold changes of signal intensity compared with the mean of the two negative control antibodies are shown. (D) Staining images showing signal colocalization between the antinucleolar antibodies and a commercial anti-Ki-67 antibody on human tonsil tissue. Scale bars indicate 50 μm.

HEp-2 reactivity was slightly more common in HLA cross-reactive antibodies (27%, 4/15 vs. 16%, 11/69), but this difference was not statistically significant (p = 0.54, chi-squared test). These results suggest that the anti-HLA reactivity of mAbs derived from intrarenal B cells is due to polyreactivity not associated with autoreactivity as defined by HEp-2 binding.

In marked contrast, when we examined mAbs from clonally expanded plasma cells from patient 6, 76 % of (16/21) antibodies had HEp-2 reactivity (Figure 7A and Supplementary Figure 6A). Reactivity was broadly distributed across eight different clonal families. These antibodies had a high prevalence of HEp-2-reactivity, especially nucleolus-reactive antibodies (Figure 7B and Supplementary Figure 5A). That multiple clones had remarkably similar patterns on staining suggested that different *in situ* clonally expanded plasma cells were targeting the same or similar antigens.

To identify potential antigens, we selected three antinuclear antibodies (6-2A4, 6-2B9 and 6-1B5) from different clonal families (Figure 7B): 6-A4 represented the most expanded clonal family. 6-2B9 showed the strongest signal in HEp-2 staining, and 6-1B5 showed the most specific nucleoli binding. Using these antibodies, we performed western blot on a nuclear fraction of HEp-2 cell lysates. Of the three antibodies tested, 6-2A4 and 6-2B9 showed similar broad immunoreactivity to antigens with relative molecular weight > 50 kDa (Supplementary Figure 6B). The 6-1B5 mAb did not detectably bind antigens.

Next, we performed immunoprecipitations (IP) with the three mAbs from the lysates of HEp-2 nuclear fractions. Two negative control mAbs were included: 7-1A5 and 7-1E3 which did not bind to either HLA or HEp-2 cells. Immunoprecipitations were washed extensively and resolved by sodium dodecyl sulfate–polyacrylamide gel electrophoresis (SDS-PAGE). Gels were Coomassie-stained and lane regions above IgG heavy chain (> 50 kDa) were excised and subjected to tandem mass spectrometry. Detected peptides were mapped to the human proteome according to UniProt database *(2017)*. Fold changes of signal intensity (normalized to median) were calculated for the three antinucleolar antibodies over the mean intensity of the two negative control antibodies.

These experiments revealed that for two of the three positive mAbs tested (6-2A4 and 6-2B9), the nucleolar antigen Ki-67 was the top hit (Figure 7C). For the other antibody, 6-1B5, another nucleolar antigen HEATR1 as the top hit. Interestingly, HEATR1 was also found in the top 10 hits of 6-2B9, and thus it is possible that these antibodies were selected for the same protein complex.

We next examined if 6-2A4 and 6-2B9 directly recognized Ki-67. As Ki-67 is large (3,256 amino acids), we expressed seven recombinant fragments in total covering the whole length of Ki-67. We expressed these fragments in E. coli, and whole lysates were resolved by SDS-PAGE and tested for antibody binding by western blot. The commercial anti-Ki67 antibody preferentially bound fragment 3 (aa 994-1489) with less binding to other fragments (Supplementary Figure 6C). In contrast, both 6-2A4 and 6-2B9 bound to fragment 6 (aa 2446-2940) with 6-2A4 also binding to fragment 7 (aa 2927-3256). These data demonstrate that both *in situ* selected antibodies directly bound similar Ki-67 domains.

Since Ki-67 was detected by two independent clones, we searched for other anti-Ki-67 antibodies by tissue staining. Given that Ki-67 is highly expressed in proliferating cells, we used tonsil tissue staining to test colocalization of the signal from the anti-nucleolar antibodies with that from a commercial anti-Ki-67 antibody. We detected a total of six antibodies that showed colocalization with Ki-67, including 6-2A4 and 6-2B9 (Figure 7D). These additional four antibodies belonged to the same clonotypes as either 6-2A4 or 6-2B9 indicating their clonal selection for Ki-67 reactivity. These results suggest that breach of self-tolerance and strong selection for self-antigens can occur in kidney of renal allografts during rejection.

Within the clonal family of 6-2A4, two antibodies (6-2H10 and 6-2G1) did not show detectable signal on tonsil staining. Therefore, we investigated their phylogenetic tree to see whether Ki-67 reactivity was gained or lost during clonal evolution. An analysis of heavy chains showed that costaining-positive 6-2G7 and negative 6-2G1 shared the same amino acid sequence (Supplementary Figure 7A). However, these two antibodies had different light chain sequences (Supplementary Figure 7B). Therefore, their different staining patterns are due to different light chain mutations. For both heavy and light chains, 6-2A4 had sequences most similar to predicted germline. These observations suggest that some mutations resulted in loss of Ki-67 binding while with other mutations it was preserved. However, across all clonally related antibodies, anti-nucleolar reactivity persisted. In toto, our results demonstrate that in allograft rejection, intrarenal B cells can breach tolerance and become strongly selected by nucleolar antigens. Furthermore, the clonal relatedness between intrarenal B cells and plasma cells indicate that local self-antigens can drive *in situ* differentiation into antibody secreting cells.

The clonal relatedness between B cells and plasma cells in patient 6 indicated that self-reactivity can occur in the local B cell pool. Furthermore, these clonally related B cell precursors expressed highly mutated antibodies suggesting that selection might occur before detectable clonal expansion. We postulated that these putative selecting antigens would be locally expressed in the inflamed kidney. Therefore, we cloned and expressed 28 representative highly mutated antibodies expressed by intrarenal B cells from patients 1-5 and 7 (4-5 antibodies per patient) (Supplementary Table 5). We then determined whether they bound inflamed allograft or normal kidney. The epitope-tagged immunoglobulin heavy chains allowed us to discern between monoclonal antibody binding and endogenous immune complexes.

Out of 28 tested antibodies, six antibodies showed detectable binding to inflamed allograft renal tissue (Supplementary Figure 8). All antibodies showed specific distributions of staining with nuclear or perinuclear binding being common. However, only 4-2C3 showed nearly ubiquitous staining with the other antibodies only binding some cell types, often tubules. These observations suggest selection by specific intrarenal expressed antigens.

Four of the six antibodies (3-1E12, 4-2E2, 5-1C3 and 7-1B2) were HEp-2-reactive with three having predominately cytoplasmic staining. However, none of the HEp-2 patterns were predictive of binding to renal tissue. This was striking for 3-1E12 which bound specifically to a subset of renal cells yet gave a diffuse cytoplasmic HEp-2 pattern, and 4-2C3 which bound renal nuclei diffusely yet was HEp-2 negative. These data indicate that the HEp-2 could be misleading in evaluating tissue specific autoantibodies. Furthermore, these data are consistent with *in situ* immunity directed against renal, and non-ubiquitous, expressed antigens.

We next examined if targeted antigens were preferentially expressed in inflamed kidney. Indeed, three antibodies (4-2E2, 5-1C3 and 7-1B2) did not bind normal kidney. However, the other three antibodies demonstrated similar binding patterns in inflamed and normal renal tissue. Therefore, there was selective binding for both inflammation-associated and normal renal antigens. Collectively, these data indicate a broad loss of tolerance in the intrarenal B cell compartment and selection by renal-associated antigens.

## Discussion

Herein, we demonstrate that in allograft rejection, intrarenal B cells have a unique gene expression profile that most closely resembles the B1 innate B cells that have been described in mice but not in humans *(Baumgarth, 2017; Glass et al., 2020)*. In contrast to murine innate B cell populations, which are constitutively present *(Baumgarth, 2017)*, Bin cells were associated with inflammation and overall transcriptional differences were most prominent in those cells expressing mutated IgG antibodies. These data suggest that in humans, B cell innate-like features are induced by specific pathways of activation and selection.

Indeed, one of the major differences between B cell populations in humans compared to mice is the lack of cells with an innate B1 cell transcriptional signature. While innate-like functions have been ascribed to human B cell subpopulations *(Rothstein and Quach, 2015)*, extensive scRNA-seq has failed to confirm their existence *(Glass et al., 2020)*. It was difficult to map Bin cells to known populations of human B cells. Rather, to characterize these cells we relied on transcriptional profiles of murine B1 innate cells. Using this approach, we identified *AHNAK* and *AHNAK*-covariant genes as a defining signature of Bin cells. While its role in B cells is unclear, AHNAK has diverse functions. For example, in mice Ahnak is required for T-cell-receptor induced intracellular calcium mobilization *(Matza et al., 2008)*. It also modulates TGF-β/Smad signaling *(Lee et al., 2014)*, which is important for various B cell functions *(Tamayo et al., 2018)*. AHNAK is likely to be functional in B cells as it is highly expressed in murine peritoneal B1 cells.

Another innate feature of Bin cells was the expression of the proinflammatory cytokine IL-15. A component of its receptor complex, IL-15RA, was also expressed in rejected renal allograft tissue. IL-15 can act directly or via trans-presentation by IL-15RA *(Dubois et al., 2002)*. Abundance of IL-15 in rejected renal allografts has been long known *(Pavlakis et al., 1996)* and antagonization of IL-15 improves graft survival. However, this is the first report to suggest that intrarenal B cells are a significant source of IL-15 in allograft rejection.

In addition to the AHNAK-gene signature, Bin cells expressed a type-I IFN signature *(Bekeredjian-Ding et al., 2005; Green et al., 2009)*. IFN-α treatment can induce rejection of renal allograft *(Magnone et al., 1995)* and the IFN-γ pathway is upregulated in rejecting renal allografts *(Saint-Mezard et al., 2009)*. IFN signaling induces TLR expression and indeed, *TLR2* and *TLR7* were upregulated in Bin cells. In mice, recipient expression of TLR2 and TLR4 is critical for renal allograft rejection *(Wang et al., 2010)*. The IFN pathway likely reflect activation mechanisms independent of the AHNAK program as we did not observe a correlation between the IFN and AHNAK signatures in single Bin cells (data not shown). Therefore, multiple activation pathways likely contribute to the molecular state of intrarenal Bin cells.

Bin cell transcriptional states were remarkably similar across patients and very different than that observed for tonsil B cells. This was particularly true of class-switched Bin cells. IgM expressing Bin cells showed some overlap with tonsil B cells and were broadly distributed in the t-SNE space. In contrast, class-switched B cells clustered away from class-switched tonsil B cells. These data suggest that, in the context of renal allograft rejection, B cell antigen selection is associated with a stereotypical and unique cell state.

Bin cells were also notable for the lack of gene signatures previously associated with renal inflammation. It is becoming clear that DN B cells are an important population in lupus nephritis that likely reflects the predominant roles of both TLR and IFN activation in disease pathogenesis *(Jenks et al., 2018)*. However, in contrast to scRNA-seq studies of intrarenal B cells in lupus nephritis, we did not observe increased expression of either *TBX21* or the DN gene signature. Hypoxia is also a feature of both human lupus and some murine lupus models *(Davidson et al.,*

*2015)*. In contrast, we did not observe upregulation of canonical hypoxia response genes. Finally, Bin cells did not express hallmarks of Breg cells including IL-10 or IL-35 (*IL12A* and *EBI3*) (Shen et al., 2014). Therefore, there were no clear and significant B cell regulatory subpopulations within the rejecting kidney. These data suggest that different renal inflammatory diseases are associated with very distinct B cell populations.

Renal allograft rejection is associated with the presence of serum DSAs. However, despite an extensive analysis, we could not identify even one Bin cell that expressed antibodies specific for HLA, be they donor specific or otherwise. This, in conjunction with a previous report of one renal explant *(Porcheray et al., 2012)*, indicate that DSAs are rarely selected for in the kidney. Rather, it is likely that DSAs are a manifestation of systemic allo-immunity associated with endovascular immune complex deposits *(Zhang, 2018)*.

Interrogation of Bin cells expressing highly mutated IgG antibodies revealed that immunoreactivity with renal tissue was relatively common. Each antibody had a different distribution of renal binding indicating specificity for different antigens. However, the restricted distributions of staining, and the poor correlations with HEp-2 staining patterns, suggest preferential binding to renal expressed, and not ubiquitous, antigens. Some antibodies bound preferentially to inflamed renal tissue. This is consistent with observations in lupus nephritis in which *in situ* selected B cells recognize vimentin which is a molecular feature of inflammation and renal tubular damage *(Kinloch et al., 2014)*. Therefore, in contrast to the diversity in B cell transcriptional states, *in situ* breaking of tolerance to local self-antigens, including patterns of inflammation and damage, might be a common feature of renal inflammation.

In the one patient in which we captured a substantial plasma cell population, we observed clonal relatedness between Bin cells and plasma cells indicating that local antigens can drive both selection and differentiation *in situ*. Furthermore, clonal expansion and ongoing somatic hypermutation, central mechanisms of antigen-driven selection, were evident. Multiple antibodies bound nucleolar antigens with two, from different clonal trees, binding directly to Ki-67. Ki-67 is associated with cell proliferation and therefore would be preferentially expressed in inflamed but not normal kidney in which the vast majority of cells are quiescent. Overall, our data is consistent with *in situ* selection for both Bin cells and plasma cells expressing antibodies reactive with antigens expressed in rejecting kidneys.

Serum autoantibodies are associated with allograft rejection. It has been postulated that many of the targeted antigens are cryptic and only exposed after ischemia-reperfusion injury *(Cardinal et al., 2017)*. It has been demonstrated that these autoantibodies can mediate vascular injury and accelerate graft rejection *(Cardinal et al., 2013)*. Many of these antibodies are thought to be natural antibodies, present prior to transplantation, titers of which predict subsequent graft loss *(Gao et al., 2014)*. In contrast, our results demonstrate that intra-renal B cells express highly mutated IgG autoantibodies which bound renal antigens. However, none of these antibodies targeted the vascular endothelium. Therefore, both natural and acquired B cell autoimmunity, targeting different antigens and compartments, are features of renal allograft rejection.

While renal reactive B cells were both present and selected in the kidney, an exhaustive characterization indicated that these cells constituted a minority of all infiltrating B cells. Most *in situ* B cells expressed unmutated or pauci-mutated IgM antibodies and therefore are unlikely to have been selected by antigen. These cells could reflect non-specific trapping of cells such as has been observed in a murine model of allograft rejection *(Walch et al., 2013)*. Another possibility is that there is preferential selection for B cells expressing low-affinity anti-renal allograft antibodies. Consistent with this, low affinity GC memory B cell precursors express CCR6^+^ B which is also expressed by intrarenal B cells *(Suan et al., 2017)*. Furthermore, as reported for CCR6^+^ B cell memory precursors, intrarenal B cells had high *S1PR1* and low *S1PR2* expression. These results suggest that intrarenal B cells might be preferentially recruited from specific peripheral B cell populations.

In addition to antigen selection, there are many potential points of interaction and cooperativity between Bin cells and inflamed allograft tissue that likely serve to retain and activate intrarenal B cells. Upregulated adhesion and migration receptor/ligand pairs include *CD55*-*CD97*, *COL4A4*-*ITGA2*, *COL1A2*-*CD44*, *ICAM2*-*ITGB2*. In addition, expressed ligand-receptors are predicted to modulate B cell activation including the receptors CD44, BTLA and ITGAM and their ligands in inflamed renal parenchyma. Finally, potential autocrine loops expressed by Bin cells, including GDF7/BMPR2 and ITCAM/ICAM2 may determine B cell activation *in situ*. The presence of both activating and inhibitory loops suggest the regulation of intrarenal B cells is complex.

In summary, we demonstrate that in renal allograft rejection, infiltrating B cells have a unique transcriptional state that suggests that they are driven by, and likely contribute to, specific innate signaling pathways and networks. Furthermore, within this population, and the plasma cells they can give rise to, are cells expressing highly mutated IgG antibodies reactive with renal expressed antigens including molecular patterns of inflammation and/or tissue damage. Based on these observations, we propose that Bin cells are not involved in alloimmune responses. Rather they function to amplify subsequent inflammatory pathways mediating end-organ damage. Because of the inflammatory networks they contribute to, and the specificity of the antibodies they express, Bin cells bridge and integrate innate and adaptive immunity to drive local inflammation.

## Materials and Methods

### Clinical sample collection

Kidney biopsies were collected as an additional biopsy core from consented patients. Presence of antibody-mediated rejection was clinically confirmed for all the sequenced transplant patients. Tonsil samples were deidentified and collected from tonsillectomy cases. All the clinical samples were collected on the day of biopsy at the University of Chicago Hospital, and approved by the Internal Review Board at the University of Chicago.

### Cell sorting

Within 5 hours after collection, tissues were minced and digested with Liberase TL (Sigma-Aldrich) for 15 minutes at 37 °C. Cells were washed and stained for 30 minutes at 4 °C with Calcein AM (eBioscience) and antibodies: PE-CD19 (eBioscience, SJ25C1), APC-CD38 (BD, HIT2), PE-Cy7-CD45 (eBioscience, HI30). Stained cells were washed, and DAPI (Thermo Fisher Scientific) was added to the single-cell suspension immediately before the samples were subjected to BD FACSAria Fusion for sorting. Doublets were excluded by FSC-A/FSC-H gating, and CD45+ Calcein+ DAPI-CD19+ CD38+ activated B cells were single-cell sorted into 96-well plates with catching buffer (RLT lysis buffer (Qiagen) with 1% 2-mercaptoethanol (Sigma-Aldrich)). Sorted cells were immediately spun down and stored at -80 °C until being processed for scRNA-seq.

### scRNA-seq

scRNA-seq was performed following Smart-seq 2 protocol *(Picelli et al., 2014)*. mRNA was purified from sorted cell lysates using SPRI beads (Beckman Coulter), and reverse transcribed to cDNA with ERCC spike-in controls (Thermo Fisher Scientific). cDNA was amplified for 20 cycles using KAPA HiFi HotStart ReadyMix PCR Kit (Kapa Biosystems). Aliquots of the amplified cDNA were also used for antibody cloning later. cDNA library was generated using Nextera XT DNA Library Preparation Kit (Illumina), pooled and sequenced with Illumina sequencer.

### Read alignment, quality control and data integration

For mapping sequencing reads, a human transcriptome (GRCh38) was obtained from Ensembl database. Low-complexity regions were masked from the transcriptome using RepeatMasker 4.1.0 *(Smit et al.) with “-noint -norna -qq” options. The masked transcriptome was used for pseudoalignment by kallisto 0.46.1 (Bray et al., 2016).* For the first cohort, poor-quality cells were excluded from the analyses if they were expressing less than 3,000 genes or more than 15,000 genes. Furthermore, to exclude cells which could be non-B cells, cells were filtered out if a sum of log-count per million (cpm) of immunoglobulin heavy chain constant region genes was below 5. Batch effects were corrected by normalizing counts to ERCC using the RUVSeq 1.16.1 *(Risso et al., 2014)* with “k = 2” option. For the second cohort, read alignment and QC was done in the same manner except that 1,000 genes were used for gene count cut off. In order to normalize the difference in sequencing depths between the first and second cohorts, SCTransform in Seurat R package 3.1.1 *(Hafemeister and Satija, 2019; Stuart et al., 2019)* was applied. The normalized data were further processed by ComBat in sva package R package 3.32.1 *(Leek et al., 2012)* to remove batch effects between the two cohorts.

### t-SNE projection and cell cluster assignment for the differential gene expression analysis

Gene expression similarity among single cells were visualized by t-SNE plots, whose coordinates were calculated by Rtsne package 0.15. Expressed Ig isotypes were determined from the scRNA-seq data by assigning the most highly expressed Ig constant region gene. Cells were categorized as “unswitched” if their isotype was IgM or IgD, and categorized as “switched” otherwise. For the first cohort, the ERCC-normalized data was scaled by log2 cpm before making t-SNE plots. For the integrated data, clusters were assigned by Seurat, and the plasma cell cluster was identified by *PRDM1* expression. Plasma cells were removed from differential expression analyses.

### Differential gene expression analysis

Differential expression was tested on genes expressed by at least 10% of each category to be compared. For the first cohort, raw pseudocounts were rounded and subjected to edgeR 3.26.8 *(Robinson et al., 2010)* with unwanted variables calculated by RUVSeq in the design matrix. For the integrated data, independent t-tests were applied to expression values after ComBat. For both analyses, false discovery rate (FDR) was calculated by adjusting p values for multiple testing by the Benjamin-Hochberg method. Genes with FDR ≤ 0.05 and log2 fold change ≥ 1 were categorized as differentially expressed.

### Hierarchical clustering of 2855 DEG

For the first cohort, differential expression was tested in four comparisons (class-switched renal vs. tonsil, unswitched renal vs. tonsil, renal class-switched vs. unswitched, and tonsil class-switched vs. unswitched). Mean expression values of identified 2855 DEG were calculated in four populations (renal switched, renal unswitched, tonsil switched, tonsil unswitched). Then hierarchical clustering was performed based on their expression pattern across the four population means, identifying six gene clusters. A heatmap was produced using pheatmap R package 1.0.12 based on the clustering and Z-scores calculated from the mean values.

### Pathway enrichment analysis

GO and KEGG enrichment was tested using clusterProfiler 3.12.0 *(Yu et al., 2012)* and org.Hs.eg.db annotation database 3.8.2. FDR 0.05 was used for significance cutoff. When there were more than 10 significantly enriched GO terms, redundant terms were removed using “simplify” function in clusterProfiler library with its default setting.

### Enrichment analysis of AHNAK-covariant genes

Gene expression in mouse B-cell subsets in the spleen or peritoneal cavity were fetched from Immgen *(Heng and Painter, 2008)*. Genes were identified as *Ahnak*-covariant genes, when their expression pattern within the peripheral B-cell subsets had a correlation coefficient ≥ 0.8 with *Ahnak*. The *Ahnak*-covariant genes were converted to their human orthologs as *AHNAK*-covariant genes using Ensembl database *(Kinsella et al., 2011)*. Then enrichment of the *AHNAK*-covariant genes in DEG clusters was tested by hypergeometric test. For the background frequency, we used the frequency of the *Ahnak*-covariant genes within all the mouse genes detected in Immgen microarray data.

### Calculation of gene expression scores

Geneset-based scores were calculated as a sum of scaled expression values of genes present in each geneset. For DEG cluster scores in mouse B-cell subsets, DEGs in each gene cluster were converted to mouse orthologs in the same manner described above. Then, scores for the mouse genes were calculated for each replicate in Immgen data. A mean score was calculated for each B-cell subset, scaled to Z-scores and visualized as a heatmap. For innate immune genes, genes tagged to “innate immune response” GO term were identified in the DEG clusters, and used to calculate a score. For ABC-associated genes, 17 ABC-upregulated genes and 9 ABC-downregulated genes were defined according to Arazi et al., 2019 *(Arazi et al., 2019)*. Then the difference between a scaled sum of expression of ABC-upregulated and downregulated genes were used as the score.

### Receptor/ligand co-expression analysis

For a geneset expressed by intrarenal B cells, we used DEG which were highly expressed in intrarenal B cells in the “class-switched renal vs. tonsil” comparison. To obtain a geneset expressed in rejected renal allografts, we referred to publicly available microarray data *(Sarwal et al., 2003)*. Since the data had three categories of rejection (acute, chronic, and drug- or infection-induced), they were grouped together as “rejection”. Then gene expression was compared between the rejection and normal allografts using t-test, in order to identify genes upregulated in rejected renal allografts. Within the identified DEG, genes encoding a receptor or a ligand were identified by crossmatching them with Functional Annotation of the Mammalian Genome (FANTOM) 5 database *(Ramilowski et al., 2015)*. Connections of identified ligands and receptors were visualized using Cytoscape 3.7.2 *(Shannon et al., 2003)*.

### Tissue staining

Paraffin-embedded formalin-fixed tissue blocks were sectioned by 3 μm thickness. Tissue sections were deparaffinized with xylene and ethanol, and subjected to antigen retrieval with 10 mM citrate buffer pH 6.0 (ThermoFisher Scientific). Tissue sections were blocked with Tris-buffered saline (TBS) containing 10 % normal donkey serum (Jackson ImmunoResearch Laboratories), and incubated with a combination of primary antibodies: rat or rabbit anti-CD19 (Invitrogen, 6OMP31 or abcam, EPR5906 respectively), rabbit anti-AHNAK (Proteintech, 16637-1-AP), mouse anti-IL15 (abcam, ab55276), and rabbit anti-Ki-67 (abcam, EPR3610). Antibody binding was detected by fluorophore-conjugated highly cross-adsorbed secondary antibodies (ThermoFisher Scientific), and nuclei were stained with Hoechst 33342 (ThermoFisher Scientific). For staining of FLAG-tagged recombinant antibodies cloned from rejection patients, rat anti-FLAG (ThermoFisher Scientific) was used as the secondary antibody, which was then detected with fluorophore-conjugated anti-rat IgG antibodies. Stained sections were mounted in ProLong^TM^ Gold Antifade Mountant (ThermoFisher Scientific) and analyzed on SP8 confocal microscopy (Leica).

### Antibody cloning and recombinant expression

*Variable regions of antibody* heavy and light chain genes were amplified from cDNA using nested PCR *(Smith et al., 2009)*. PCR products were Sanger-sequenced and mapped to IMGT reference. Results of heavy-chain genes were analyzed for clonality and mutation frequency. Clonal families were defined by VDJ gene usage and CDR3 length. Next, heavy and light chain variable regions were cloned into an IgG expression vector. The vector had a FLAG tag at the C terminus of IgG constant region to enable tissue staining in the presence of IgG-expressing cells or IgG deposition *(Kinloch et al.)*. A pair of cloned heavy and light chain vectors were transfected to HEK293 cells, and expressed IgG was purified using Protein A agarose beads (Pierce), eluted in 0.1 M glycine-HCl pH 2.8, neutralized with 1M Tris buffer pH 9.0 and stored in PBS with 0.05% sodium azide.

### HLA-binding assay

Antibodies were diluted at 150 μg/mL in PBS and tested on LAB Screen Mixed (OneLambda) according to the manufacturer’s protocol. To test the binding in the presence of blocking, positive antibodies from intrarenal B cells were retested in the presence of human serum proteins. Antibodies were prepared in PBS, then diluted 1:1 in PBS or negative control serum included in the kit, and subjected to the assay. NBG ratio was calculated as the experimental readout according to the manufacturer’s protocol:

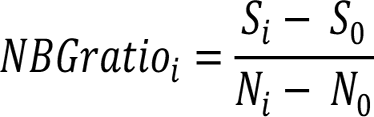

Si and S0 are sample signals from the i th antigen-coated beads and negative beads, and Ni and N0 are negative serum signals from the i th antigen-coated beads and negative beads. NBG ratio ≥ 2.2 was used as the positivity threshold. For SAB assay, differences in trimmed MFI between antibodies and negative control serum were used as a readout. Eplets information was fetched from HLA Epitope Registry *(Duquesnoy et al., 2013)*,

### HEp-2 cell staining

Antibodies were diluted at 50 μg/mL in PBS, and tested on NOVA Lite^®^ HEp-2 ANA kit (Inova Diagnostics) according to the manufacturer’s protocol. Antibody binding was detected on SP8 confocal microscopy by fluorescent signal from fluorescein isothiocyanate (FITC)-conjugated polyclonal anti-human IgG secondary antibody included in the kit.

### Western blot

HEp-2 cells (ATCC^®^ CCL-23TM) were cultured and harvested. A nuclear fraction was prepared using Nuclear Extraction Kit (Abcam) following the manufacturer’s protocol. Before the final centrifugation step, lysates were sonicated with three times of a 10-second pulse on ice. Lysates were boiled in Laemmli buffer at 95 °C for 5 minutes, and resolved by SDS-PAGE. Proteins on the gel were transferred to a polyvinylidene fluoride membrane, blocked by 5 % bovine serum albumin-containing TBS, and incubated with 10 μg/mL antibody diluted in the blocking buffer at 4 °C overnight. The membrane was washed and incubated with a horseradish peroxidase-conjugated anti-human IgG antibody, and binding was detected with Pierce™ ECL Western Blotting Substrate (ThermoFisher Scientific).

### Expression of Ki-67 fragments

Seven fragments were designed to cover the whole Ki-67 protein. Fragments 1 and 2 were cloned by PCR from cDNA of HEp-2 cells. DNA encoding the other fragments were purchased from Bio Basic Inc. Primers used and DNA fragments ordered are listed in Supplementary Table 7. DNA encoding each fragment were digested with NdeI and XhoI restriction enzymes (NEB) and cloned into pET-24b (+) vector. Rosetta (DE3) E. coli (Millipore) were transformed with the expression vectors. Overnight culture was diluted in 1:5 in LB media, and IPTG was inoculated. After 4 hours of culture, 1 mL of culture was centrifuged, and cell pellet was resuspended in RIPA lysis buffer. The lysate was used for western blot.

### IP from a nuclear fraction of HEp-2 cell lysates

HEp-2 nuclear lysates were precleared with Protein A Agarose beads, and incubated with 5 μg of antibodies at 4 °C overnight. The beads were washed with Tween20-containing PBS, and captured antibody-antigen complexes were eluted and resolved by SDS-PAGE. Gels were stained with InstantBlue^®^ Protein Stain (Expedeon) at 4 °C overnight. Stained gels were destained in deionized water, excised leaving molecular weight higher than IgG heavy chain, and used for mass spectrometry.

### Sample preparation for mass spectrometry

Gel Samples were excised by sterile razor blade and chopped into ∼1 mm^3^ pieces. Each section was washed in distilled water and destained using 100 mM NH_4_HCO_3_ pH7.5 in 50 % acetonitrile. A reduction step was performed by addition of 100 μL 50 mM NH_4_HCO_3_ pH 7.5 and 10 μL of 200 mM tris (2-carboxyethyl) phosphine HCl at 37 °C for 30 minutes. The proteins were alkylated by addition of 100 μL of 50 mM iodoacetamide prepared fresh in 50 mM NH_4_HCO_3_ pH 7.5 buffer, and allowed to react in the dark at 20 °C for 30 minutes. Gel sections were washed in water, then acetonitrile, and vacuum dried. Trypsin digestion was carried out overnight at 37 °C with 1:50 - 1:100 enzyme-protein ratio of sequencing grade-modified trypsin (Promega) in 50 mM NH_4_HCO_3_ pH 7.5, and 20 mM CaCl2. Peptides were extracted first with 5 % formic acid, then with 75 % ACN : 5 % formic acid, combined and vacuum dried. Digested peptides were cleaned up on a C18 column (Pierce), speed vacuumed and sent for liquid chromatography–tandem mass spectrometry (LC-MS/MS) to the Proteomics Core at Mayo Clinic.

### High-performance liquid chromatography (HPLC) for mass spectrometry

All samples were resuspended in Burdick & Jackson HPLC-grade water containing 0.2 % formic acid (Fluka), 0.1% TFA (Pierce), and 0.002% Zwittergent 3-16 (Calbiochem), a sulfobetaine detergent that contributes the following distinct peaks at the end of chromatograms: MH^+^ at 392, and in-source dimer [2M + H^+^] at 783, and some minor impurities of Zwittergent 3-12 seen as MH^+^ at 336. The peptide samples were loaded to a 0.25 μL C8 OptiPak trapping cartridge custom-packed with Michrom Magic (Optimize Technologies) C8, washed, then switched in-line with a 20 cm by 75 μm C18 packed spray tip nano column packed with Michrom Magic C18AQ, for a 2-step gradient. Mobile phase A was water/acetonitrile/formic acid (98/2/0.2) and mobile phase B was acetonitrile/isopropanol/water/formic acid (80/10/10/0.2). Using a flow rate of 350 nL/min, a 90-minute 2-step LC gradient was run from 5 % B to 50 % B in 60 minutes, followed by 50 % - 95 % B over the next 10 minutes, hold 10 minutes at 95 % B, back to starting conditions and re-equilibrated.

### LC-MS/MS data acquisition and analysis

The samples were analyzed via data-dependent electrospray tandem mass spectrometry (LC-MS/MS) on a Thermo Q-Exactive Orbitrap mass spectrometer, using a 70,000 RP survey scan in profile mode, m/z 360-2000 Da, with lockmasses, followed by 20 HCD fragmentation scans at 17,500 resolution on doubly and triply charged precursors. Single charged ions were excluded, and ions selected for MS/MS were placed on an exclusion list for 60 seconds.

All LC-MS/MS *.raw Data files were analyzed with MaxQuant version 1.5.2.8, searching against the UniProt Human database (Download 9/16/2019 with isoforms, 192928 entries) *.fasta sequence, using the following criteria: LFQ was selected for Quantitation with a minimum of 1 high confidence peptide to assign LFQ Intensities. Trypsin was selected as the protease with maximum missing cleavage set to 2. Carbamidomethyl (C) was selected as a fixed modification. Variable modifications were set to Oxidization (M), Formylation (N-term), Deamidation (NQ). Orbitrap mass spectrometer was selected using an MS error of 20 ppm and a MS/MS error of 0.5 Da. 1% FDR cutoff was selected for peptide, protein, and site identifications. Ratios were reported based on the LFQ Intensities of protein peak areas determined by MaxQuant (version 1.5.2.8) and reported in the proteinGroups.txt. The proteingroups.txt file was processed in Perseus (version 1.6.7). Proteins were removed from this results file if they were flagged by MaxQuant as “Contaminants”, “Reverse” or “Only identified by site”. Three biological replicates were performed. Samples were filtered to require hits to have been seen in at least two replicates per condition. Intensities were normalized by median intensity within each sample. Then, log2 fold changes over the means of negative controls were obtained for the three antinucleolar antibodies.

The full proteomic data set will be uploaded to the ProteomeXchange repository (https://www.ebi.ac.uk/pride/archive/).

## Supplementary Materials

**Supplementary Table 1. Patient information.** For transplant patients, age, gender and kidney survival length are masked to protect patients’ confidentiality. Tonsil donors are deidentified.

**Supplementary Table 2. Ig class distribution across patients.** Number of QC-passed cells grouped by Ig class are shown for each patient.

**Supplementary Table 3. A list of 2855 DEG.** Gene symbols are shown with their clusters identified in Figure 2D. Columns “comparison1-4” show which category expressed the genes significantly higher in each differential expression test.

**Supplementary Table 4. A list of Ahnak and AHNAK-covariant genes.** Gene symbols of Ahnak-covariant mouse genes and their human orthologs (*AHNAK*-covariant genes) are shown.

**Supplementary Table 5. A list of recombinantly expressed antibodies.** Information of heavy and light chain genes and assay results are shown.

**Supplementary Table 6. Eplet analysis of SAB assay-positive antibodies.** Top-10 hit antigens, trimmed mean MFI, and eplets present in the antigens are shown for each antibody. Eplets shared among all top-10 hits are colored in red.

**Supplementary Table 7. DNA sequences of primers and Ki-67 fragments.** Sequences of cloning primers and purchased DNA encoding Ki-67 fragments.

**Supplementary Figure 1.**
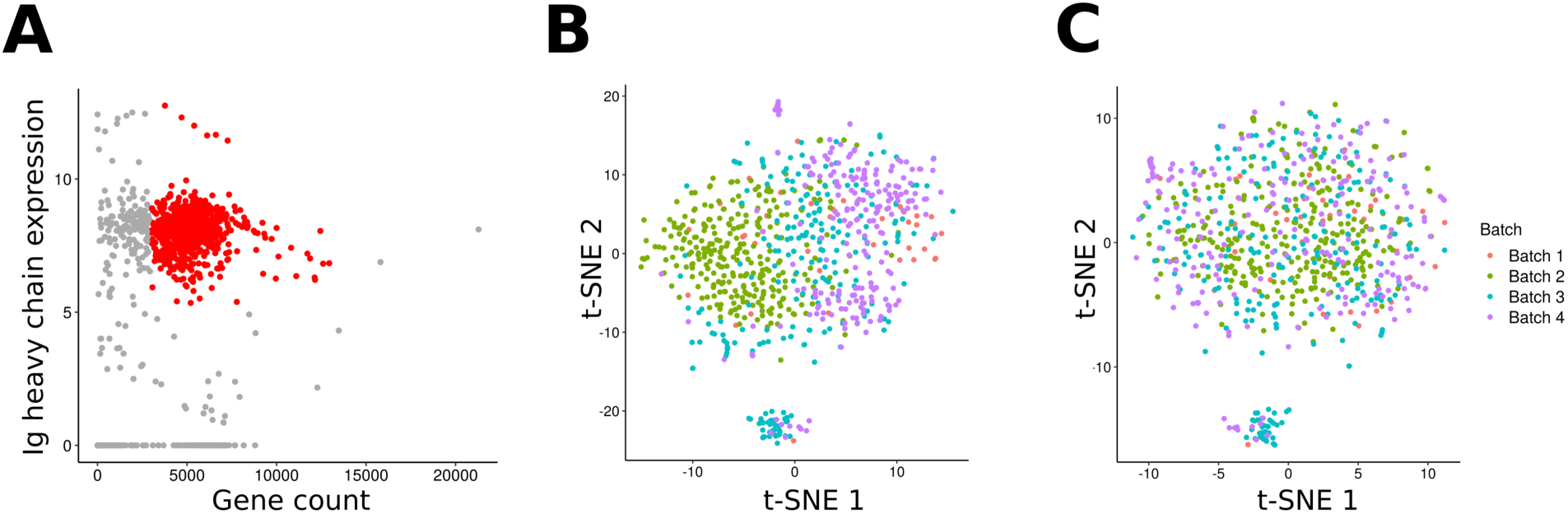
QC and normalization with ERCC spike-in to dissolve batch-dependent clusters. (A) A scatter plot showing detected gene count and Ig heavy chain gene expression on the x and y axis respectively. Cells which passed QC are colored in red. (B and C) t-SNE plots colored by experimental batches before (B) and after (C) batch normalization with ERCC spike-in and RUVSeq.

**Supplementary Figure 2.**
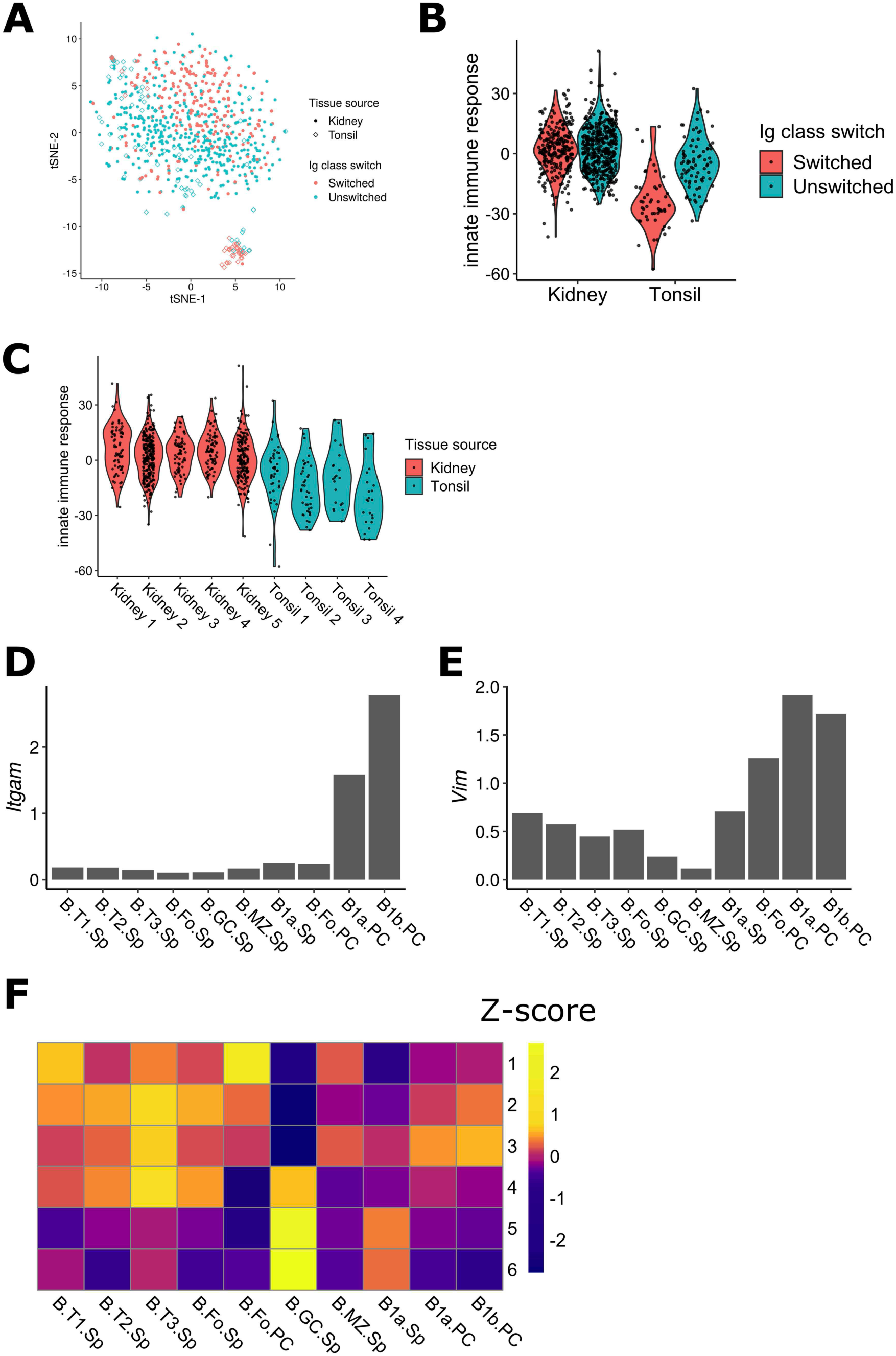
Innate immune and B1-like gene expression in intrarenal B cells. (A) A t-SNE plot as in Figure 2A, in which Ig constant region genes were removed from the data. (B-C) A sum of scaled expression values of genes tagged to “innate immune response” GO term in cluster 2-3. Data were grouped by tissue sources (B) or patients (C). (D and E) Expression of *Itgam* (D) and *Vim* (E) in the mouse B cell subsets. (F) A heatmap showing DEG scores as in Figure 3G, which was calculated without the *Ahnak*-covariant genes.

**Supplementary Figure 3.**
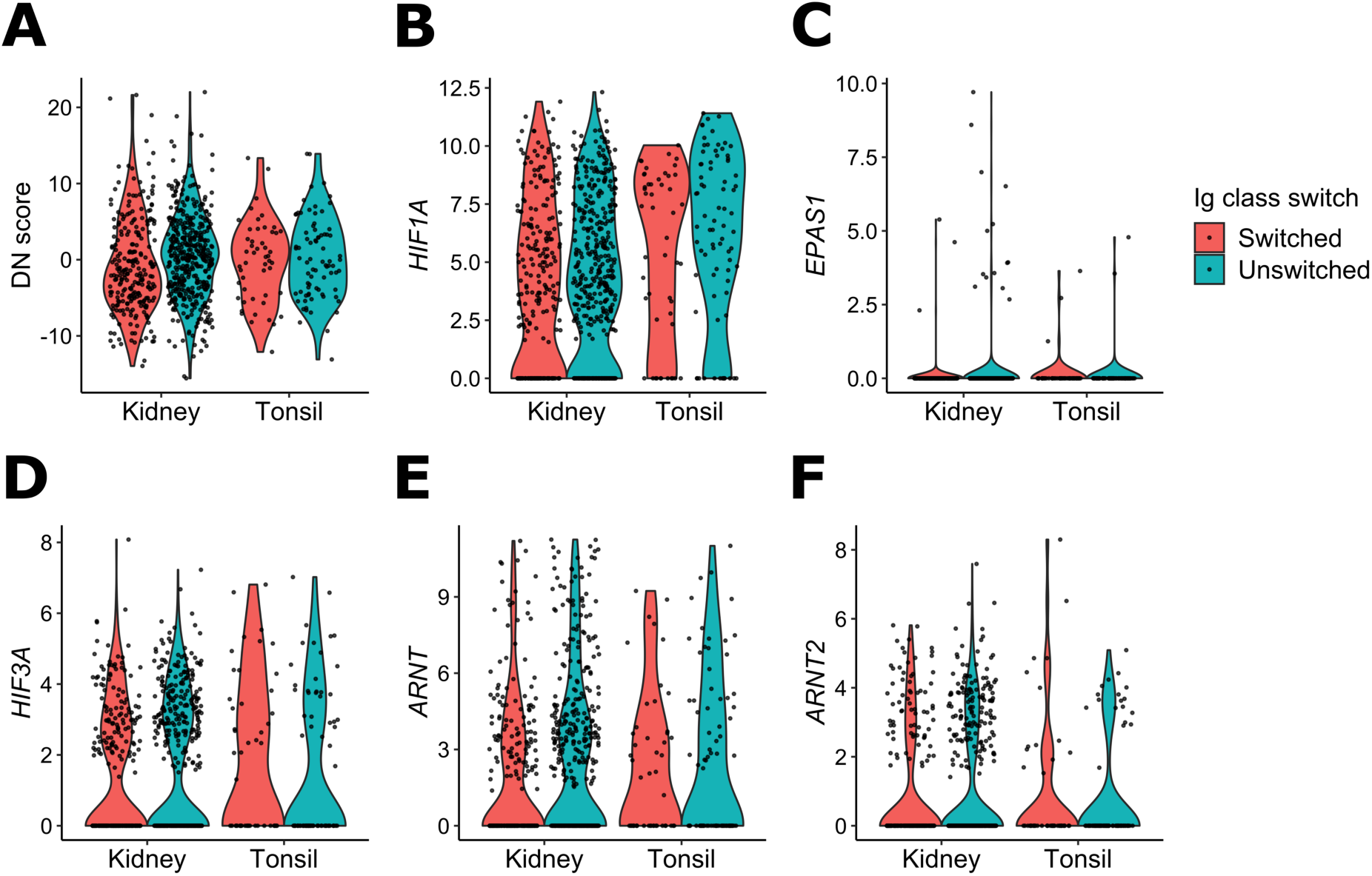
Intrarenal B cells had no association with ABC and HIF signatures. (A-F) Violin plots showing the gene expression of ABC-associated genes (A), *HIF1A* (B), *HIF2A* (C), *HIF3A* (D), *HIF1B* (E), or *HIF2B* (F). Cells were grouped by their tissue source and Ig class-switch state.

**Supplementary Figure 4.**
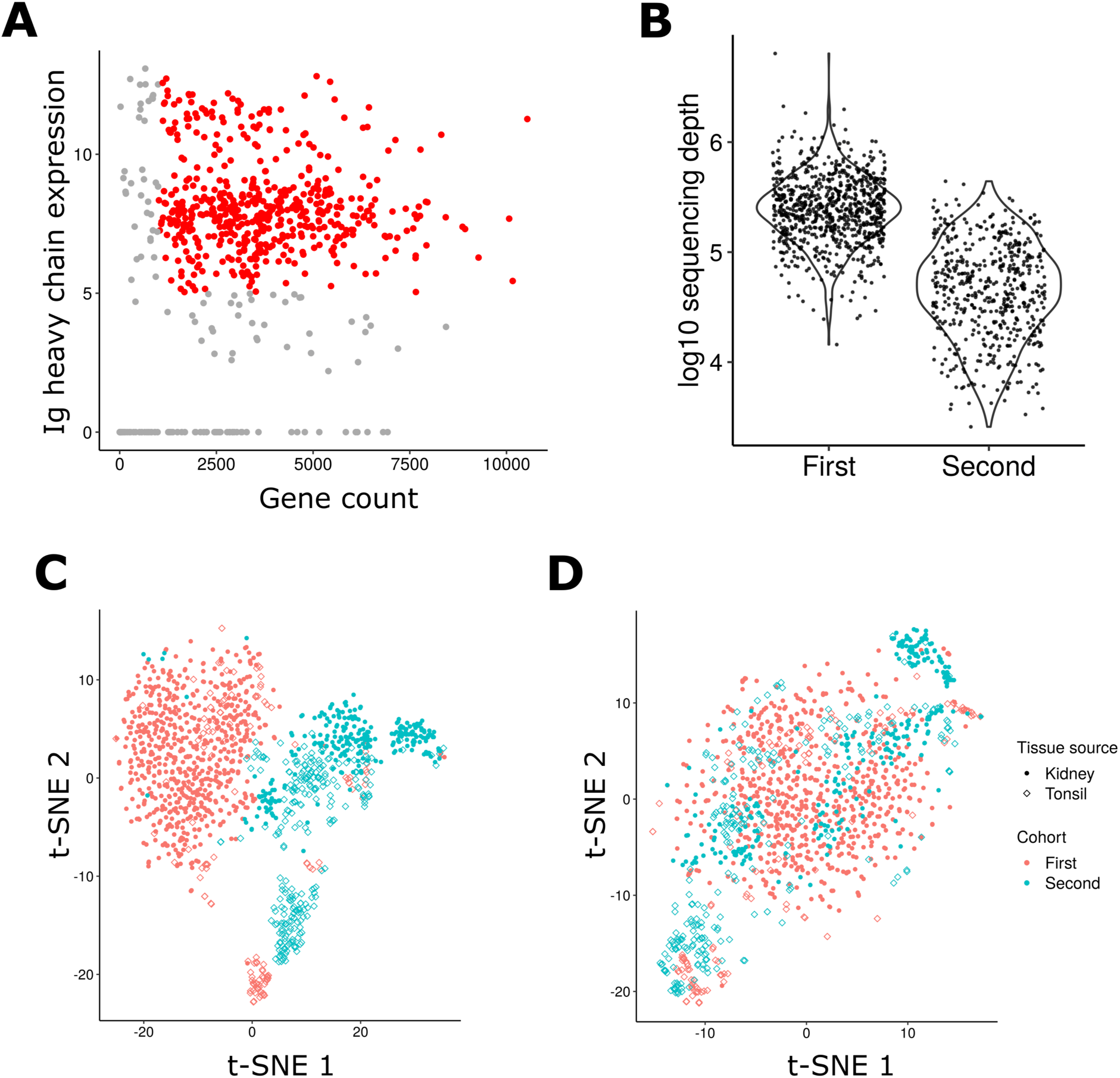
QC of the second cohort and data integration. (A) A scatter plot showing detected gene count and Ig Heavy chain gene expression on the x and y axis respectively. Cells which passed QC are colored in red. (B) A violin plot showing sequencing depth of cells which passed QC in each cohort. (C and D) t-SNE plots made from the data normalized by log2 cpm without data integration (C) or the data normalized by SCTransform and integrated by ComBat (D). Color and shape respectively indicate cohorts and tissue source.

**Supplementary Figure 5.**
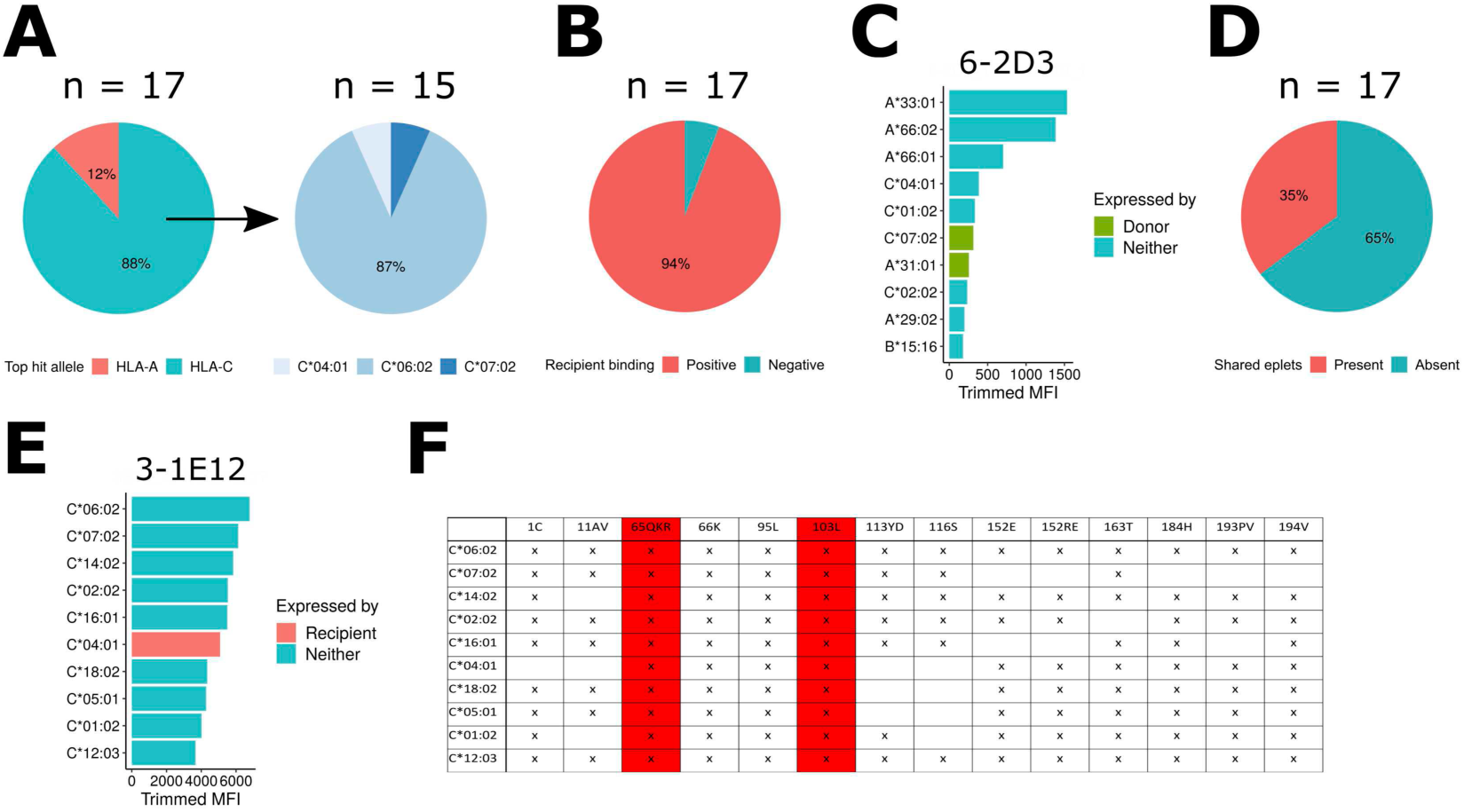
SAB assay analysis. (A) Pie charts showing top-hit alleles in the SAB assay. 17 antibodies with trimmed MFI ≥ 1,000 were analyzed. (B) A ratio of antibodies that had a recipient-expressed antigen among top-10 hits. (C) Trimmed MFI of 6-2D3. Donor-expressed antigens are colored in green. (D) A ratio of antibodies whose top-10 hits shared eplets. (E) Trimmed MFI of a representative antibody 3-1E12 with a shared eplet. A recipient-expressed antigen is colored in red. (F) Eplets (columns) present in antigens (rows) detected in (E). Eplets shared among more than five antigens are shown. Eplets shared among all the top-10 hits are highlighted in red.

**Supplementary Figure 6.**
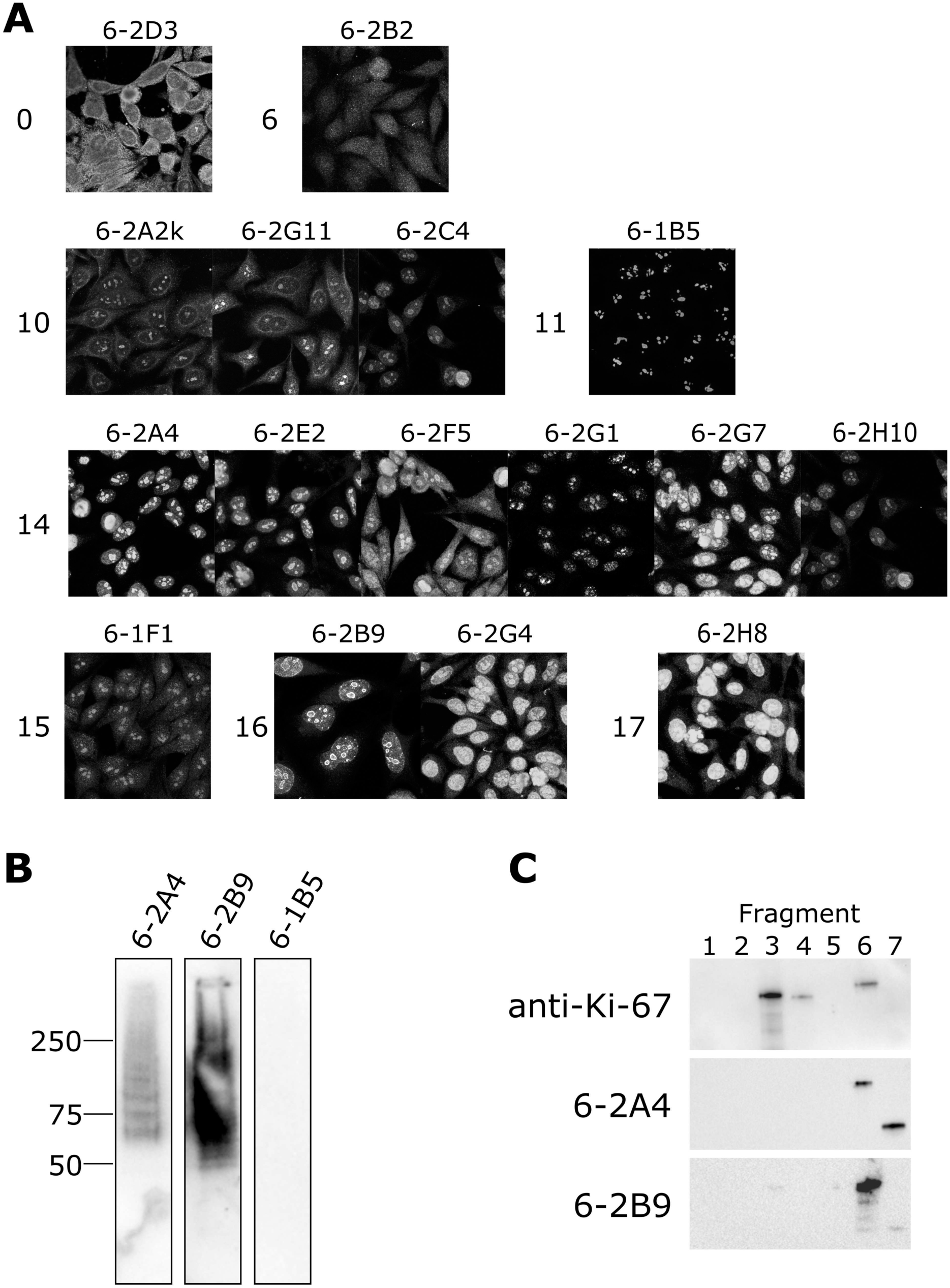
Autoreactivity of antibodies expressed by plasma cells in patient 6. (A) HEp-2 staining images of the 16 positive antibodies cloned from patient 6 plasma cells. Antibodies of the same clonal family are grouped together. Numbers on the left side of the images indicate their clonal family number as in Figure 6C. (B) Western blot on HEp-2 nuclear lysates using the three antibodies (6-2A4, 6-2B9, 6-1B5). (C) Western blot on E. coli lysates expressing Ki-67 fragments. Each fragment contains these amino acids: fragment 1, aa 1-526; fragment 2, aa 509-1009; fragment 3, aa 994-1489; fragment 4, aa1476-1976; fragment 5, aa1963-2459; fragment 6, aa2446-2940; and fragment 7, aa2927-aa3256.

**Supplementary Figure 7.**
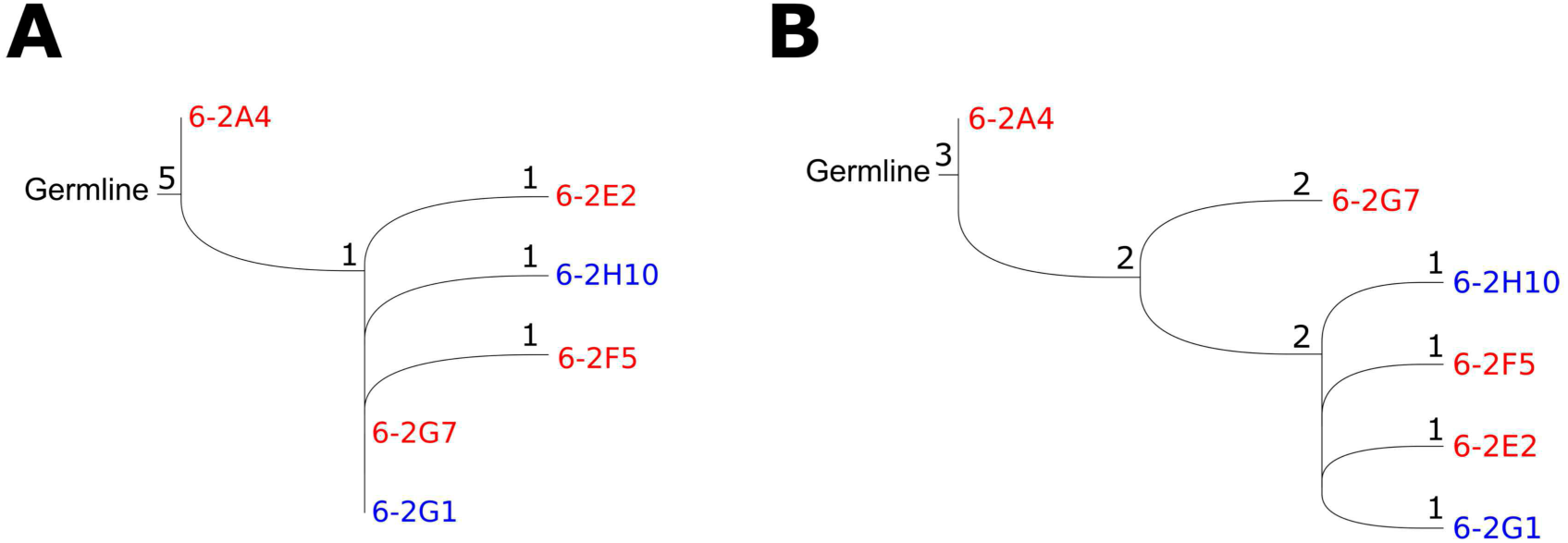
Changes in Ki-67 reactivity through clonal evolution. (A and B) Phylogenetic trees showing amino acid changes in heavy (A) and light (B) chains of antibodies in clonal family 14 that 6-2A4 belongs to. Numbers by nodes indicate amino acid differences from their previous nodes. Color indicates Ki-67 costaining results (red: positive, blue: negative).

**Supplementary Figure 8.**
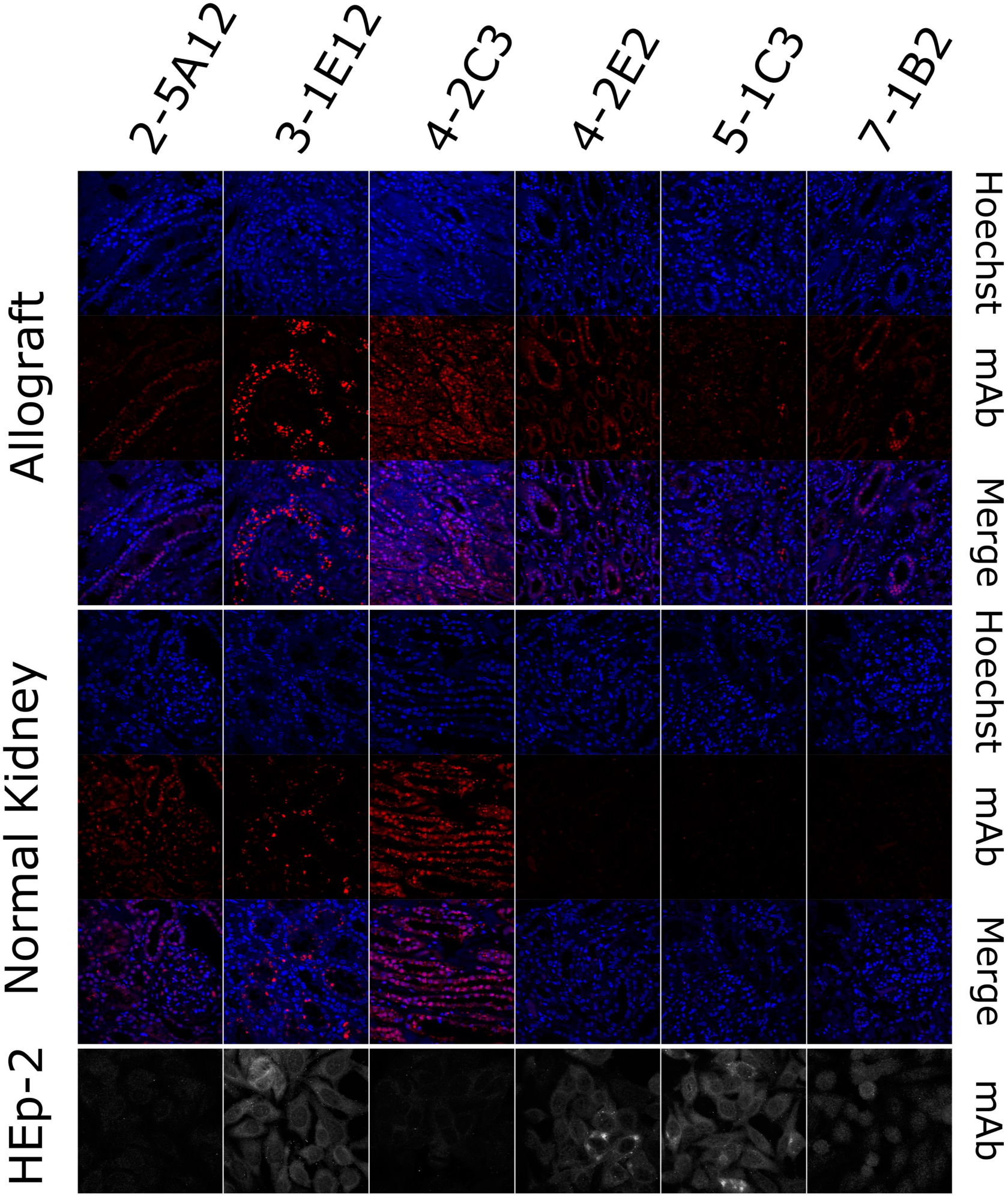
Bin cells express IgG antibodies that bind renal antigens. Indicated Flag-tagged antibodies were used to probe inflamed allograft or normal renal tissue. Antibodies were also assayed for HEp-2 immunoreactivity. Images are representative (n=2).

